# A ‘How-to’ Guide for Interpreting Parameters in Habitat-Selection Analyses

**DOI:** 10.1101/2020.11.12.379834

**Authors:** John Fieberg, Johannes Signer, Brian Smith, Tal Avgar

## Abstract

1. Habitat-selection analyses allow researchers to link animals to their environment via habitat-selection or step-selection functions, and are commonly used to address questions related to wildlife management and conservation efforts. Habitat-selection analyses that incorporate movement characteristics, referred to as *integrated step-selection analyses*, are particularly appealing because they allow modeling of both movement and habitat-selection processes.
2. Despite their popularity, many users struggle with interpreting parameters in habitat-selection and step-selection functions. Integrated step-selection analyses also require several additional steps to translate model parameters into a full-fledged movement model, and the mathematics supporting this approach can be challenging for many to understand.
3. Using simple examples, we demonstrate how weighted distribution theory and the inhomogeneous Poisson point-process can facilitate parameter interpretation in habitat-selection analyses. Further, we provide a “how to” guide illustrating the steps required to implement integrated step-selection analyses using the amt package.
4. By providing clear examples with open-source code, we hope to make habitat-selection analyses more understandable and accessible to end users.

## Introduction

New technologies (e.g., improved Global Positioning System [GPS] collars) and advances in remote sensing have made it possible to collect animal location data on unprecedented spatial and temporal scales (Kays, Crofoot, Jetz, & Wikelski, 2015; Robinson et al., 2020), which in turn has fueled the development of new methods for modeling animal movement and for linking individuals to their environments (Guisan, Thuiller, & Zimmermann, 2017; Hooten, Johnson, McClintock, & Morales, 2017). Two of the most popular approaches for analyzing telemetry data, *habitat-selection functions* (HSFs; Box 1) and *step-selection functions* (SSFs), compare environmental covariates at locations visited by an animal (“used locations”) to environmental covariates at a set of locations assumed available to the animal (“available locations”) using logistic and conditional logistic regression, respectively (Boyce & McDonald, 1999; Fortin et al., 2005; Thurfjell, Ciuti, & Boyce, 2014). These methods are widely available in most statistical software packages, and thus, they provide a robust and easy-to-implement framework for analyzing habitat-selection patterns. Note, here and throughout, we use the term *habitat-selection function* rather than the traditional *resource-selection function* to highlight our broader interest in modeling the effects of a diverse set of environmental variables (e.g., those capturing risks and environmental conditions in addition to resources). Habitat-selection functions are used to identify habitat features that are preferentially used or avoided by a species, and thus, to infer ecological needs and limitations, generate expected distribution maps, and inform demographic projections across space and time in support of species and landscape management (Boyce & McDonald, 1999; Matthiopoulos et al., 2015; Matthiopoulos, Field, & MacLeod, 2019).

Step-selection functions are further used to identify fine-scale behavioral interactions between animals and their biotic and abiotic environment (e.g., Dickie, McNay, Sutherland, Cody, & Avgar, 2020). Despite their popularity, our collective experience has been that many users struggle to interpret parameters in HSFs and SSFs. Further, it seems that papers attempting to address this issue have had limited success, and in some aspects may have increased confusion (see e.g., Keating & Cherry, 2004; Johnson, Nielsen, Merrill, McDonald, & Boyce, 2006; Lele, Merrill, Keim, & Boyce, 2013; Avgar, Lele, Keim, & Boyce, 2017; Chamaille-Jammes, 2019).

Here, we highlight how point-process models and weighted distribution theory provide simple and effective frameworks for interpreting regression parameters in habitat-selection and step-selection functions. In the sections that follow, we begin by reviewing recent research connecting habitat-selection functions to point-process models and weighted distribution theory. Using these connections, we demonstrate correct interpretation of parameters using simple examples of models fit to GPS locations of fisher (*Pekania pennanti*) from upstate New York (LaPoint et al., 2013a, 2013b). We then provide a short review of step-selection functions, including their history and methods for parameter estimation. Step-selection analyses (Box 2) are particularly appealing because: 1) they provide an objective method for defining habitat availability in terms of movement constraints; they relax the assumption that locations are statistically independent; and 3) by including movement characteristics (e.g., functions of step length and turn angle) as predictors, they provide a means to model both movement and habitat-selection processes (termed an *integrated step-selection analysis* by Avgar, Potts, Lewis, & Boyce, 2016). Recognizing that many may find the mathematics supporting integrated step-selection analyses intimidating, we aim to provide a “how to” guide demonstrating the steps required to implement the approach using the amt package (Signer, Fieberg, & Avgar, 2019). This demonstration is expanded upon using coded examples in the supplementary appendices, which we encourage the reader to explore. We end with a short discussion highlighting challenges related to statistical dependencies and model transferability.

## Habitat-Selection Functions (HSFs)

### Logistic Regression

Much of the confusion surrounding the interpretation of parameters in habitat-selection functions can be attributed to the use of logistic regression to model *use-availability* data (Keating & Cherry, 2004). Logistic regression is most easily understood as a model for binary random variables that can take on one of two values (0 or 1) with probability that depends on one or more explanatory variables (Hosmer, Lemeshow, & Sturdivant, 2013).

Consider, for example, a study designed to infer how various environmental characteristics influence whether a habitat patch (e.g., a contiguous area of forest) will be used by one or more animals. In this case, we may randomly select *n* habitat patches and monitor them to determine if they are used (*y*_*i*_ = 1) or not (*y*_*i*_ = 0) for *i* = 1, 2,…, *n*. Logistic regression allows us to model the probability that each patch will be used, *P*(*y*_*i*_ = 1) = *p*_*i*_, as a logit-linear function of *k* patch-level predictors (*X*_*i*1_,…,*X*_*ik*_) and regression parameters (*β*_0_*, β*_1_,…,*β*xs_*k*_):

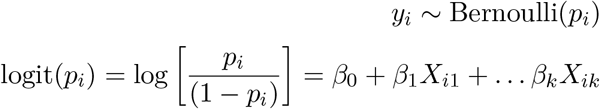

After having fit a model, we can exponentiate the regression coefficients, exp(*β*_*j*_) for (*j* = 1*,…, k*), to quantify how the odds of patch *i* being used, *p*_*i*_/(1− *p*_*i*_), change as we increase the *j*^th^ predictor by 1 unit while holding all other predictors constant. We can also use the inverse-logit transformation (eqn. (1)) to estimate the probability that patch *i* will be used, given its set of spatial predictors:

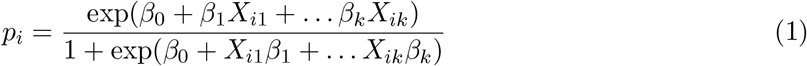

The logit transformation ensures that *p*_*i*_ will be constrained between 0 and 1 for all values of the predictor variables.

Contrast this approach with how logistic regression is used to study habitat selection. In a typical habitat-selection sample units, usually points; these groups are not mutually exclusive (i.e., available habitat may also be used). In this case, *y*_*i*_ is no longer a Bernoulli random variable since *p*_*i*_ depends on the ratio of used to available points (which is under control of the analyst). That is, the probability that a location will be a “used point” decreases with the number of user-generated “available” locations. Further, despite the fact that most analyses of telemetry data quantify environmental covariates in discrete space (i.e., pixels in a raster), the sampling itself is point-level and in continuous space. Thus, it is perhaps not surprising that there has been considerable confusion and controversy surrounding the use of logistic regression with use-availability data (e.g., Keating & Cherry, 2004; Johnson et al., 2006; Chamaille-Jammes, 2019).

Various arguments have been constructed to justify the use of logistic regression when analyzing use-availability data (Manly, McDonald, Thomas, McDonald, & Erickson, 2002; Johnson et al., 2006; Aarts, MacKenzie, McConnell, Fedak, & Matthiopoulos, 2008), but a significant breakthrough came when Warton & Shepherd (2010) made a connection between logistic-regression and a spatial inhomogeneous Poisson point-process (IPP). A spatial IPP is a model for random locations in space, where the expected spatial density of the locations depends on spatial predictors (see next section, **Inhomogeneous Poisson Point-process Model**). Warton & Shepherd (2010) showed that as the number of available points is increased towards infinity, the slope parameters in logistic regression models will converge to the slope parameters in an IPP model. Interestingly, several other popular approaches for analyzing species distribution data, including MaxEnt (Phillips & Dudík, 2008; Elith et al., 2011), weighted distribution theory with an exponential form (Lele & Keim, 2006), and resource utilization functions (Millspaugh et al., 2006), have been shown to be equivalent to fitting a spatial IPP model (Warton & Shepherd, 2010; Aarts, Fieberg, & Matthiopoulos, 2012; Fithian & Hastie, 2013; Hooten, Hanks, Johnson, & Alldredge, 2013; Renner et al., 2015).

Instead of focusing on *p*_*i*_, as is typical in applications to presence-absence data, logistic regression applied to use-availability data should simply be viewed as a convenient tool for estimating coefficients in a *habitat-selection function*, *w*(*X*(*s*); *β*) = exp(*X*_1_(*s*)*β*_1_ + … *X*_*k*_(*s*)*β*_*k*_) (Boyce & McDonald, 1999; Boyce, Vernier, Nielsen, & Schmiegelow, 2002), where we have written *X*(*s*) to highlight that the predictors correspond to measurements at specific point locations in geographic space, *s*. As we will see in the next section, this expression is equivalent to the intensity function of an IPP model but with the intercept (the log of the baseline intensity) removed; the baseline intensity gives the expected density of points when all covariates are 0. Because habitat-selection functions do not include this baseline intensity, they are said to measure “relative probabilities of use”, or alternatively, said to be “proportional to the probability of use” (Manly et al., 2002). Although the term *probability of use* sounds appealing, probability in continuous space can only be assigned to areas, not points. Further, although probability of use is easily defined for discrete sample units (e.g. grid cells), these probabilities should increase with the size of the spatial unit and also with the study duration (Lele & Keim, 2006; Lele et al., 2013). Thus, with telemetry studies, it seems more natural to model spatial (or spatio-temporal) intensity functions or rates of use in continuous space (and time). Subsequently, “probabilities of use” can be determined by integrating these intensity functions over whatever spatial (and temporal) unit is deemed appropriate. Point-process models allow us to do just that.

### Inhomogeneous Poisson Point-Process (IPP) Model

The IPP model provides a simple framework for modeling the density of points in space as a log-linear function of spatial predictors through a spatially-varying intensity function, *λ*(*s*):

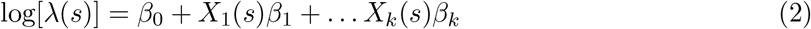

where *s* is a location in geographic space, and *X*_1_(*s*), …, *X*_*k*_(*s*) are *k* spatial predictors associated with location *s*. The intercept, *β*_0_, determines the log-density of points (within a small homogeneous area around *s*) when all *X*_*j*_(*s*) (*j* = 1, …, *k*) are 0, and the slopes, *β*_1_, …, *β*_*k*_, describe the effect of spatial covariates on the log density of points in space. The IPP model can be understood by listing its key features and assumptions, namely:

1. The number of points in an area *G*, *y*_*G*_, is a Poisson random variable with mean *E*[*y*_*G*_] = *∫*_*G*_ *λ*(*s*)*ds* (the spatial integral of *λ*(*s*) over *G*).
2. Locations are independent (any clustering can be explained by spatial covariates).

If all available spatial predictors are measured only at a coarse scale (e.g., at a set of gridded or rasterized cells), then fitting the IPP model is equivalent to fitting a Poisson regression model (Aarts et al., 2012). Specifically, one may treat the counts, *y*_*i*_, in *n* discrete spatial units (*i* = 1, …, *n*), as a set of independent Poisson random variables with means = *λ*(*s*_*i*_)|*G*_*i*_| where *λ*(*s*_*i*_) is given by eqn. (2) and |*G*_*i*_| is the area of unit *i*. Note that log[*E*(*y*_*i*_)] = log[*λ*(*s*_*i*_)|*G*_*i*_|] = log[*λ*(*s*_*i*_)] + log(|*G*_*i*_|). Thus, the log-link used in Poisson regression implies the area, |*G*_*i*_|, should be included as an offset (a predictor variable with regression coefficient fixed at a value of 1).

When spatial predictors are available at the point-level, as will be the case whenever constructing “distance to” predictors (e.g., distance to nearest road, water source, etc), it will be advantageous to model the locations in continuous space. In telemetry studies, the density of points will be determined by the frequency and duration of data collection. Thus, *β*_0_ will not be of biological interest, and it will be appropriate to focus efforts on estimating and interpreting the slope coefficients, *β*_1_, …, *β*_*k*_, which determine relationships between the spatial covariates and the relative density of locations throughout the study area (Fithian & Hastie, 2013). As is the case with linear and generalized linear models (e.g., Poisson regression), we can estimate parameters using maximum likelihood or Bayesian methods. Both approaches require writing down an expression, called the *likelihood*, that captures the data-generating mechanism in terms of one or more parameters. With telemetry data, it makes sense to work with the conditional likelihood of the IPP model (Aarts et al., 2012), i.e., the likelihood of the observed locations in space, conditional on there being *y*_*G*_ total observed locations. The conditional likelihood is given by:

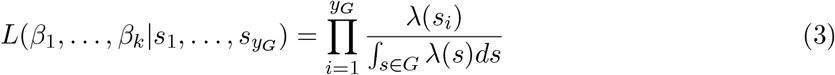

where the product is over the *y*_*G*_ observed locations, *λ*(*s*_*i*_) is the intensity function evaluated at observation *i*, and the integral in the denominator evaluates the intensity function over the spatial domain of interest (Cressie, 1992; Aarts et al., 2012). If we plug *λ*(*s*_*i*_) = exp(*β*_0_ + *X*_1_(*s*_*i*_)*β*_1_ + *… X*_*k*_(*s*_*i*_)*β*_*k*_) into eqn. (3), *β*_0_ will cancel from the numerator and denominator, leaving us with:

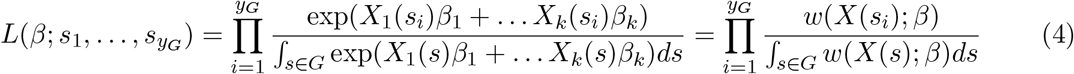

where *w*(*X*(*s*); *β*) = exp(*β*_1_*X*_1_(*s*) + *… β*_*k*_*X*_*k*_(*s*)) is our habitat-selection function.

The binomial likelihood associated with logistic regression differs from eqn. (4), but Warton & Shepherd (2010) showed that logistic regression estimators of slope coefficients converge to the those of the IPP model as the number of available points increases toward infinity. Thus, the connection to the IPP model addresses a common question that arises when estimating habitat-selection functions, namely, “how many available points do I need?” The exact answer depends on how difficult it is to estimate the integral in the denominator of eqn. (4); the recommendation we offer is to increase the number of available points until the estimated slope coefficients no longer change much. Fithian & Hastie (2013) later showed that the convergence results of Warton & Shepherd (2010) hold only if the model is correctly specified, but assigning “infinite weights” to available points ensures the results hold more generally. Therefore, when fitting logistic regression or other binary response models (e.g., boosted regression trees) to use-availability data, we also suggest assigning a large weight (say 5000 or more) to each available location and a weight of 1 to all observed locations (larger weights can be used to verify that results are robust to this choice). For a coded example in R (R Core Team, 2019), see section **Interpreting Parameters in Habitat-Selection Functions** and Supplementary Appendix A.

### Weighted Distributions

Weighted distribution theory provides another way to interpret parameters in habitat-selection functions (Lele & Keim, 2006; Johnson, Thomas, Ver Hoef, & Christ, 2008). Let:

- *u*(*X*) = the frequency distribution of habitat covariates, *X*, at locations used by our study animals.
- *a*(*X*) = the frequency distribution of habitat covariates, *X*, at locations assumed to be available to our study animals.

We can think of the *habitat-selection* function, *w*(*X*; *β*), as providing a set of weights that takes us from the distribution of available habitat to the distribution of used habitat:

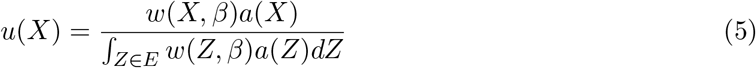

The denominator of eqn. (5) ensures that the right-hand side integrates to 1, and thus, *u*(*X*) is a proper probability distribution; the variable *Z* here is just a dummy variable used to allow integration over the frequency distribution of our environmental covariates. Because these distributions are written in terms of the habitat covariates, *X*, instead of geographical locations, we say that model is parameterized in *environmental space* (*E*) (Hirzel & Le Lay, 2008; Elith & Leathwick, 2009; Matthiopoulos et al., 2020b).

To show that weighted distribution theory is consistent with the IPP formulation discussed above, we can rewrite eqn. (5) in *geographic space* (*G*):

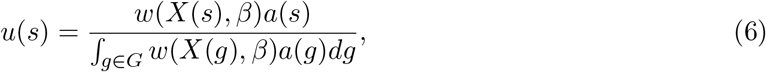

where the denominator integrates over a geographic area, *G*, that is assumed to be available to the animal and *g* is a dummy variable for integration. Here *u*(*s*) is equivalent to the utilization distribution encountered in the literature on probabilistic estimators of animal home ranges (Van Winkle, 1975; Worton, 1989; Signer & Fieberg, 2020) and tells us how likely we are to find an individual at location *s* in geographic space. The utilization distribution, *u*(*s*), depends on the environmental covariates associated with location *s*, through *w*(*X*(*s*); *β*), and the distribution of available locations in geographic space, *a*(*s*). When fitting HSFs, *a*(*s*) is typically assumed to be a uniform distribution within the geographical domain of availability, *G* (e.g., the individual’s home range, the population’s range, or the species range depending on the hierarchical level of habitat selection of interest; Johnson, 1980), with all areas within *G* assumed to be equally available to the organism. Hence, *a*(*s*) is typically a constant, 1/|*G*|, that cancels from the numerator and denominator. Then, if we let *w*(*X*(*s*); *β*) = exp(*β*_1_*X*_1_(*s*) + *… β*_*k*_*X*_*k*_(*s*)), we end up with the conditional likelihood of the IPP model (eqn. (4)) (Aarts et al., 2012). In summary, the IPP model and weighted distribution theory with an exponential form provide equivalent, suitable frameworks for interpreting parameters in logistic regression models fit to use-availability data.

### Interpreting Parameters in Habitat-Selection Functions

To demonstrate how the IPP and weighted distribution theory frameworks help with interpreting parameters in fitted habitat-selection functions, we now consider a simple example using 3,004 locations of a fisher named Lupe tracked as part of a larger telemetry study (LaPoint et al., 2013a, 2013b). These data are publicly available and have been featured in a workshop highlighting Movebank’s *Env-DATA* system for annotating locations with environmental covariates (Dodge et al., 2013; Fieberg et al., 2018). The location data were combined with available points sampled randomly from within a minimum convex polygon (MCP) formed using Lupe’s locations. The used and available locations were then transformed to a projected coordinate reference system (NAD83 / Conus Albers) and annotated with environmental variables measuring human population density (Center for International Earth Science Information Network (CIESIN) Columbia University & CIAT, Centro Internacional de Agricultura Tropical, 2005), elevation (U. S. / Japan ASTER Science Team, 2009), and landcover class (Defourny et al., 2009). The original landcover data were grouped to form a variable named `landuseC` with the following categories: `forest, grass` and `wet` (Fig. 1). We created centered (mean = 0) and scaled (SD = 1) variables labeled `elevation` and `popden` from the original elevation and population density variables. We also created an indicator variable, `case_`, taking on a value of 1 for all used points and 0 for all available points (later, we discuss how to choose the number of available points).

**Figure 1:**
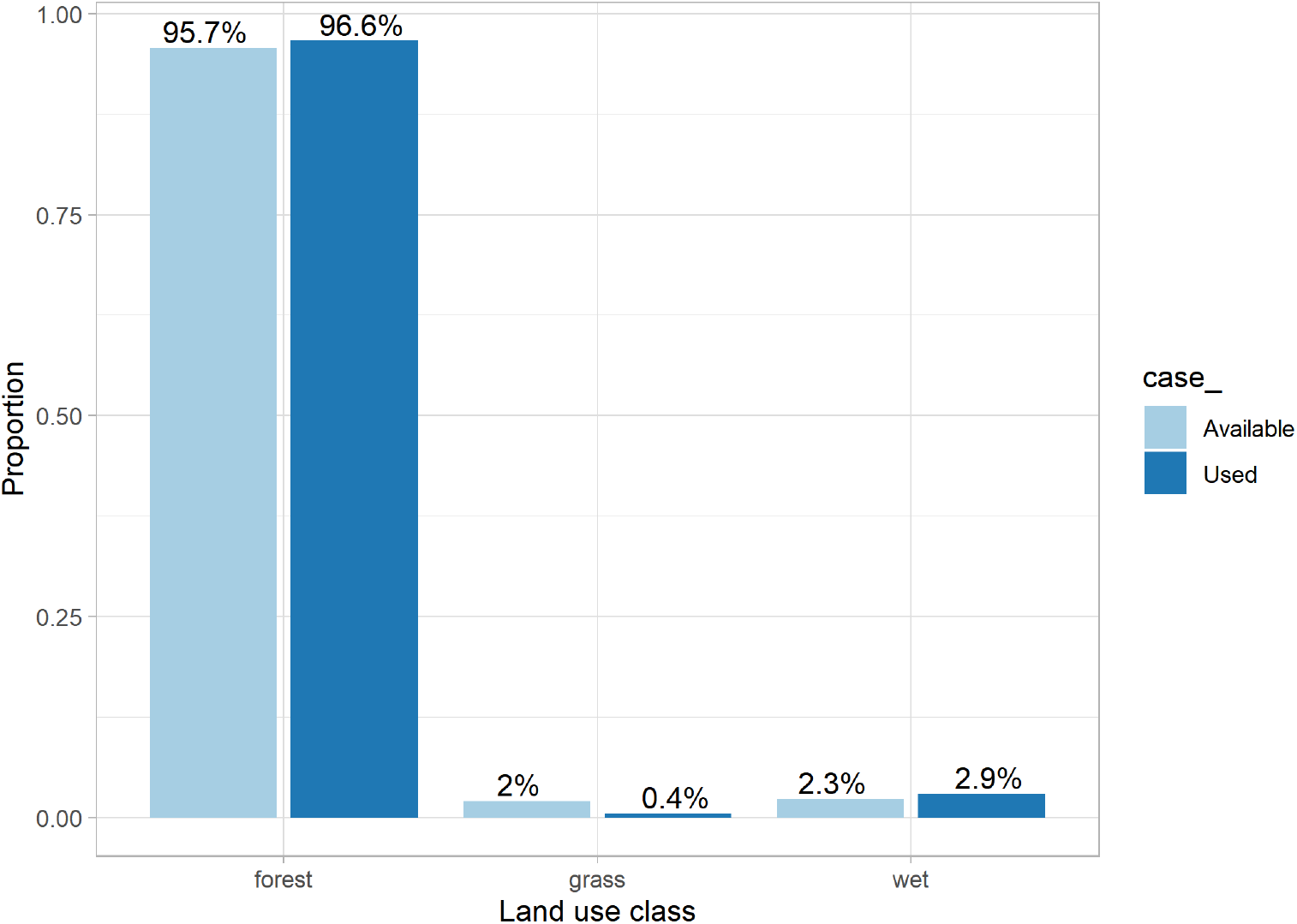
Distribution of used and available locations among different landscape cover classes for a fisher in upstate New York (LaPoint et al., 2013a, 2013b).

For ease of interpretation, we will begin by assuming the effects of elevation, population density, and landcover class are additive and linear (on the log scale; eqn. (2)). Later, we will discuss how we can relax these assumptions using interactions to allow the effect of covariates to depend on the value of other habitat covariates and polynomials or splines to to relax the assumption of linearity. We assign a weight of 5000 to the available locations and a weight of 1 to all observed locations (Fithian & Hastie, 2013). We can then fit a weighted logistic regression model using the `glm` function in R:

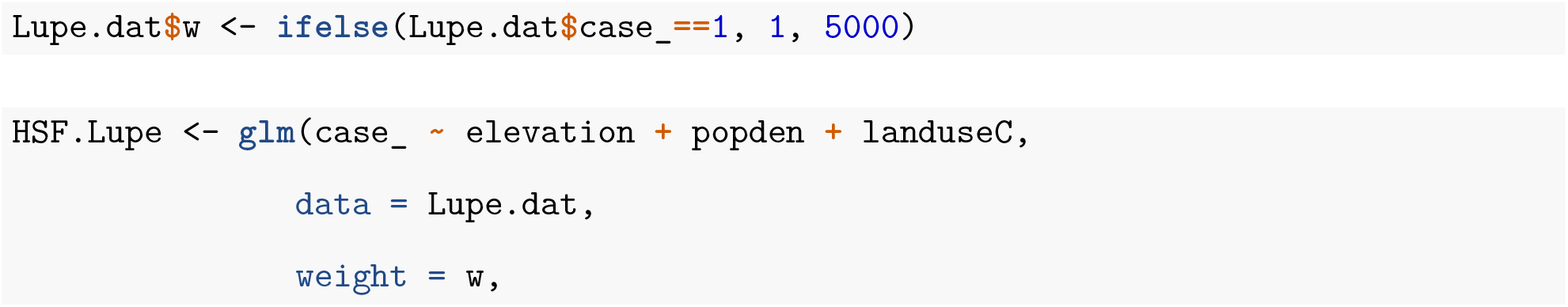
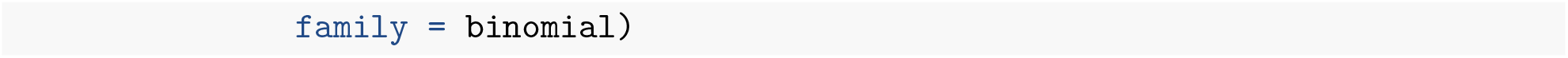

Before interpreting the coefficients, it is important to make sure we have included a sufficient number of available points to allow parameter estimates to converge to stable values. To evaluate parameter stability, we fit logistic regression models to data sets with increasing numbers of available points (from 1 available point per used point to 100 available points per used point; see Supplementary Appendix A for the code). The intercept decreased as we increased the number of available points (as it is roughly proportional to the log difference between the numbers of used and available points), but the slope parameter estimates, on average, did not change much once we included at least 10 available points per used point (Fig. 2). Further, as expected, estimates varied less from sample to sample as we increased the number of available points. Thus, we conclude that, in this particular case, having 10 available points per used point is sufficient for interpreting the slope coefficients. The only downside to including even more available points is that it may slow down computations, which is not an issue here. Increasing the number of available points also further reduces Monte Carlo error, so we proceed with the largest sample size we explored (100 available points per used point).

**Figure 2:**
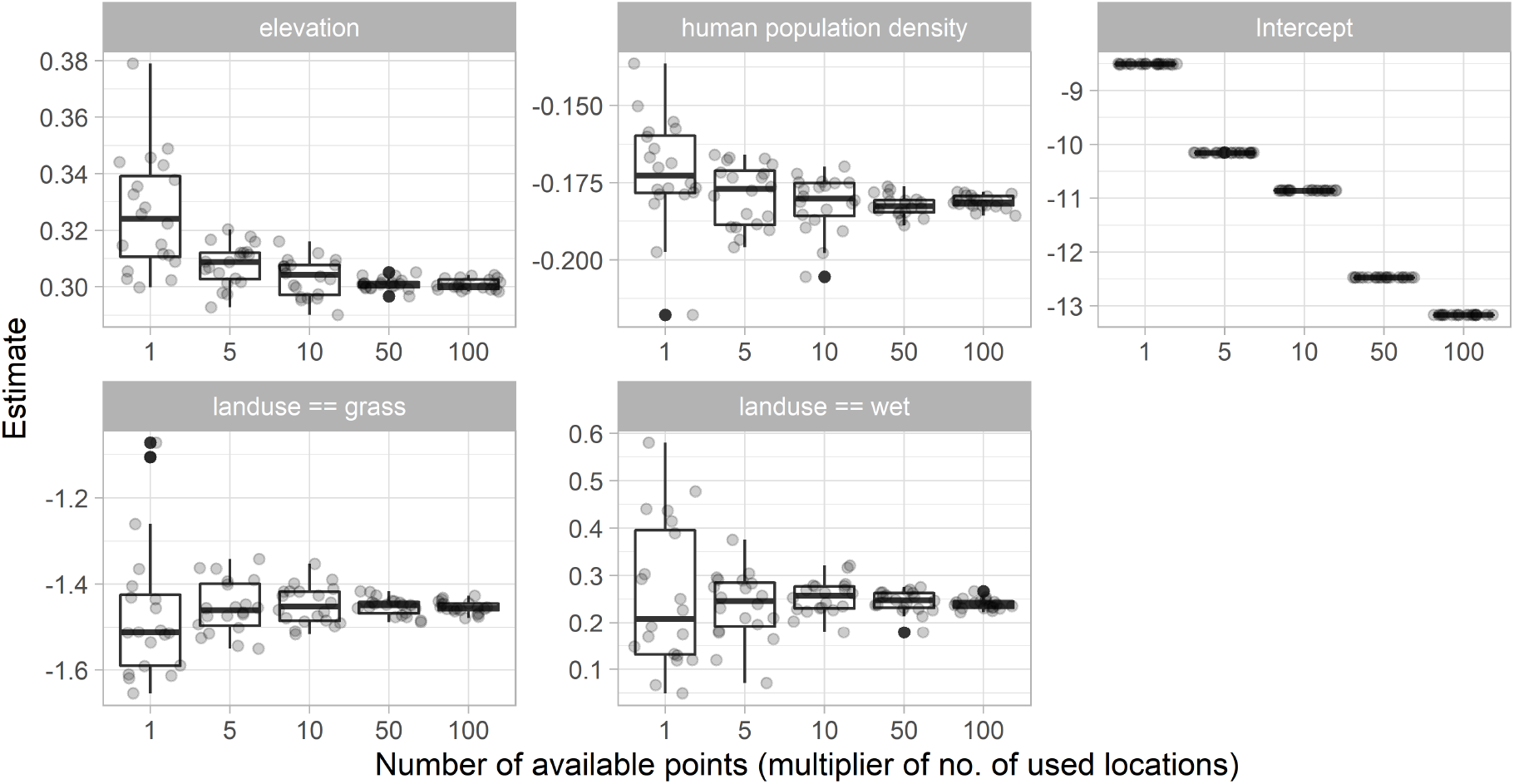
Estimated parameters in fitted habitat-selection functions using increasing numbers of available points. Each dot represents an estimate from fitting a logistic regression model to 3004 GPS telemetry locations combined with a random sample of available points, with sample size given by the x-axis (where 1 means 3004 available points and 100 means 300,400 available points).

Let’s consider the interpretation of the continuous covariates reflecting elevation and population density (Table 1, *Model 1*). Qualitatively, we might infer from the positive coefficient for `elevation` and negative coefficient for `popden` that, all other things being equal, Lupe is likely to select locations at higher elevations and in areas of lower population density. But, how do we interpret these coefficients quantitatively? Consider the following two locations, both in the same landcover class and with the same associated population density, but differing by 1 unit in `elevation` (since we have scaled this variable, a difference of 1 implies that the two observations differ by 1 SD in the original units of elevation):

- location *s*_1_: `elevation` = 3, `popden`=1.5, `landuseC` = wet
- location *s*_2_: `elevation` = 2, `popden`=1.5, `landuseC` = wet

**Table 1:** Regression coefficients (SE) in fitted habitat-selection functions fit to data from Lupe the fisher. Models 1 and 3 use forest as the reference level, Model 2 uses wet as the reference level. Model 3 includes interactions between elevation and landcover classes.

|  | Model 1 | Model 2 | Model 3 |
| --- | --- | --- | --- |
| (Intercept) | -13.168<br>(0.019) | -12.918<br>(0.107) | -13.171<br>(0.020) |
| elevation | 0.303<br>(0.017) | 0.303<br>(0.017) | 0.313<br>(0.017) |
| popden | -0.183<br>(0.021) | -0.183<br>(0.021) | -0.186<br>(0.021) |
| landuseCgrass | -1.477<br>(0.278) |  | -1.471<br>(0.278) |
| landuseCwet | 0.250<br>(0.108) |  | 0.183<br>(0.116) |
| landuseC1forest |  | -0.250<br>(0.108) |  |
| landuseC1grass |  | -1.727<br>(0.297) |  |
| elevation:landuseCgrass |  |  | 0.112<br>(0.380) |
| elevation:landuseCwet |  |  | -0.498<br>(0.127) |

Using eqn. (6), we can calculate Lupe’s relative use of location 1 versus location 2:

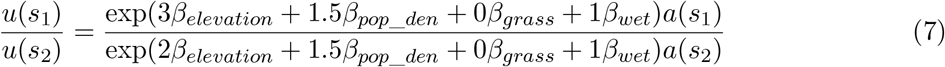

where we have dropped the integral from eqn. (6) because it appears in both the numerator and denominator (and thus, cancels out). Now, if both locations are *equally available*, then *a*(*s*_1_) = *a*(*s*_2_), and we have:

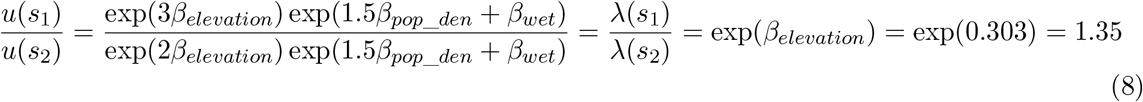

Thus, we see that this ratio also provides an estimate of the relative intensity of use of the two locations (i.e., *λ*(*s*_1_)*/λ*(*s*_2_)), assuming the locations are equally available. In the context of habitat-selection analyses, Avgar et al. (2017) refer to exp(*β*) as quantifying *relative selection strength* (RSS).

Note that we would arrive at the exact same expression if we chose *any* two locations that differed by 1 unit of `elevation` and had the same values for `popden` and `landuseC`. Thus, exp(*β*_*elevation*_) quantifies the relative intensity of use of two locations that differ by 1 SD unit of `elevation` but are otherwise equivalent (i.e., they are equally available and have the same values of all other habitat covariates). If Lupe were to be presented with two such hypothetical locations, the model suggests she would be 1.35 times more likely to choose the one with the higher elevation. A similar interpretation can be ascribed to `popden`. Given two observations that differ by 1 SD unit of `popden` but are otherwise equal, Lupe would be exp(−0.183) = 0.833 times as likely to choose the location with higher population density (or, equivalently, exp(0.183) = 1.20 times more likely to choose the location with the lower population density).

What about the coefficients for the landcover categories? Looking again at the regression output (Table 1, *Model 1*), we see that `grass` has a negative coefficient and `wet` has a positive coefficient. It is tempting to infer that Lupe spends most of her time in wet areas and rarely spends time in grassy habitats. As Figure 1 makes it clear, however, these inferences are not exactly correct. First, it is important to understand how categorical predictors are encoded in regression models. There are a number of different ways to parameterize the effect of categorical variables and unfamiliar readers may want to work through an introductory regression text (e.g., Chapter 6 of Kéry, 2010). The default coding in R is to treat one of the levels (whichever comes first alphanumerically) as a reference level and then to create a set of dummy variables that contrast the remaining levels of the categorical variable with this reference level. In our case, `forest` is the reference level. The coefficients associated with `grass` and `wet` represent contrasts between these land cover classes and the `forest` class. Qualitatively, we can use the signs and absolute magnitude of the coefficients for `grass` and `wet` to rank the landcover classes in terms of their relative selection strength, with `grass` < `forest` < `wet`. But again, how should we interpret the coefficients for `grass` and `wet` quantitatively?

Let’s again consider 2 locations, this time assuming they have the same elevation and population densities, but with one location in wet and the other location in `forest`:

- location *s*_1_: `elevation` = 2, `popden`=1.5, `landuseC` = wet
- location *s*_2_: `elevation` = 2, `popden`=1.5, `landuseC` = forest

Lupe’s relative use of location 1 relative to location 2 is given by (eqn. (6)):

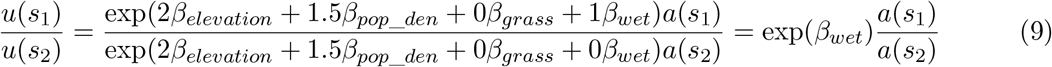

Thus, *assuming the two locations are equally available*, we might infer that Lupe would be exp(0.250) = 1.28 times more likely to choose the `wet` location than the location in `forest`. Of course, we know from Figure 1 that `forest` and `wet` are not equally available on the landscape. The higher availability of `forest` habitat implies that Lupe is more likely to be in `forest` than `wet`. We could attempt to correct for differences in availability within the MCP surrounding Lupe’s locations by multiplying our result by the ratio of habitat availability for wet relative to `forest` habitats (2.3% versus 95.7%; Fig. 1). This gives us an adjusted ratio equal to exp(0.250)(0.023)/(0.957) = 0.03, suggesting we are (1/0.03) = 33 times more likely to find Lupe in `forest` than `wet` habitat. With this calcualtion, we had to assume, perhaps naively, that the availability distributions for `popden` and `elevation` were the same in both `wet` and `forest` cover classes. In reality, if Lupe decides to move from `forest` to `wet`, it is likely that she will experience a change in `elevation` and `popden` too (i.e., these factors will not be held constant). To quantify Lupe’s relative use of `forest` versus `wet` habitat, while also accounting for the effects other environmental characteristics that are associated with these habitat types, we can use integrated intensities – i.e., we can integrate the spatial utilization distribution, *u*(*s*), over all `forest` and `wet` habitats:

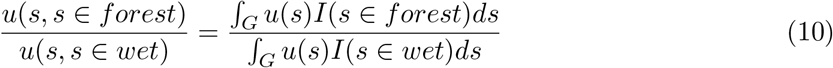

where *I*(*s* ∈ *forest*) and *I*(*s* ∈ *wet*) are indicator functions equal to 1 when location *s* is in `forest` or `wet`, respectively (and 0 otherwise). We can estimate this ratio using estimated HSF values, 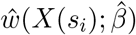, at our set of *n*_*a*_ available points drawn from within *G*. Specifically, we sum the HSF values at all available points that fall in `forest` and them divide by the sum of HSF values for all available points falling in `wet`:

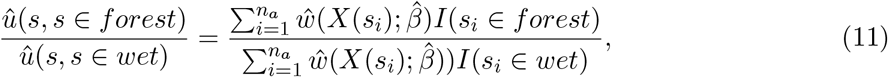

This ratio is also equal to 33, which agrees with the observed data; Lupe was found in `forest` habitat 33 times more often than in `wet` habitat (see Supplementary Appendix A for code demonstrating how to calculate these quantities in R). Thus, we conclude Lupe is 33 times more likely to be found in `forest` than `wet` habitat (despite preferring `wet` over `forest`), assuming she restricts her movements to the MCP surrounding her observed locations and all of this MCP is equally available to her.

Before moving on, it is important to note that naively-adjusted ratios (multiplying by availability of `wet` and `forest` habitats) and integrated-intensities will not always agree. In fact, we find that they differ when comparing Lupe’s relative use of `wet` versus grass habitat, with the integrated-intensity better agreeing with the observed data (see Supplementary Appendix A). Somewhat related, Avgar et al. (2017) suggested calculating *average effects* for continuous predictors, *X*, by comparing the change in relative intensities from increasing *X* by 1 unit (to *X* = *x* + 1) to the average value of *w*(*X*(*s*); *β*) for all locations *s* with *X*(*s*) = *x*. These average effects will also be influenced by cross-correlations among predictor variables included in the model.

Instead of integrating *u*(*s*) over discrete cover types, we could integrate over specific geographic areas. For example, we could use integrated intensities to compare two areas in space, replacing the “landcover class” indicator variables, *I*(*s*_*i*_ ∈ *forest*) and *I*(*s*_*i*_ ∈ *wet*) in eqn. (11), with indicator variables for whether available locations fall in particular spatial regions. In addition, we could choose to change the area of interest (and thus, area of integration) from *G* to 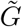, and then use the fitted model and eqn. (6) to project how Lupe would spend her time in a novel environment (referred to as an “out-of-sample” prediction). Despite the common reliance on HSFs as predictive models, out-of-sample predictions often suffer from poor accuracy, especially when compared to “in sample” predictions, i.e., predictions for the same area and time frame from which the original data were collected (Torres et al., 2015; Yates et al., 2018). We return to this important point in the discussion section.

Let’s next consider what happens if we change the reference level of the land cover variable from `forest` to `wet` (Table 1, *Model 2*).

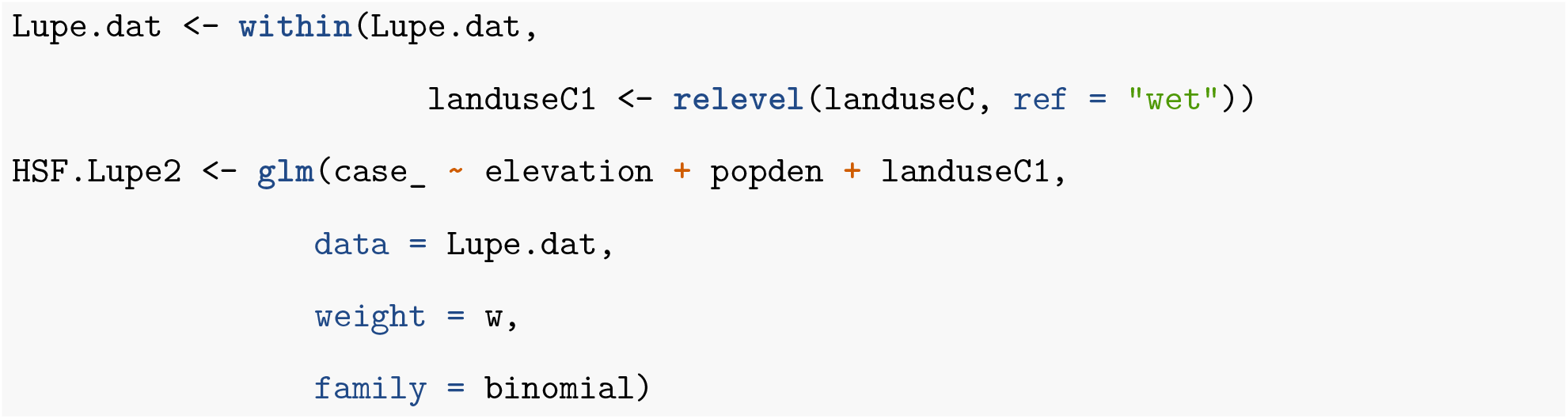

The coefficients for `elevation` and `popden` do not change. Note, however, that the coefficient for `forest` is negative despite Lupe using `forest` more than its availability (i.e., *u*(*s, s* ∈ *forest*) *> a*(*s, s* ∈ *forest*)) and Lupe spending more than 95% of her time in the forest! What is going on? Remember, the coefficients for categorical predictors reflect use:availability ratios for each level of the predictor relative to the use:availability ratio for the reference class. The coefficient for `forest` is negative because the use:availability ratio for `forest` is less than the use:availability ratio for the reference class, `wet` (see Fig. 1). Depending on the reference level, it is possible to have a positive (negative) coefficient even when that landcover class is used more (less) than its availability. Furthermore, it is possible for a landcover class to be used frequently but have a negative coefficient.

We have seen many ecologists, including some that are very quantitatively skilled and familiar with habitat-selection models, make mistakes when interpreting coefficients associated with categorical predictors. This example also highlights the importance of plotting your data (e.g., Fig. 1) and considering habitat availability when interpreting regression coefficients. Plotting distributions of covariates for both used and available locations is one of the best ways to understand fitted habitat-selection models, and is a good strategy to use for both continuous and categorical predictors (Merow, Smith, & Silander, 2013; Fieberg, Forester, et al., 2018).

### Interactions Between Environmental Predictors

Consider the distribution of `elevation` at used and available locations across the different habitat classes (Fig. 3). We see that there is a wider range of `elevation` in `forest` and `wet` habitat compared to grass habitat, and there is a clear association between `elevation` and `landuseC`, with higher median `elevation` at used locations in `forest` and grass habitat relative to `wet` habitat. Perhaps more importantly, we also see that values of `elevation` are higher, on average, for used locations (compared to available locations) in `forest` and grass, whereas the opposite is true in `wet` habitat. Although we should be skeptical of interactions that we discover while exploring our data (i.e., interactions that were not specified *a priori*), an analyst may be tempted to include an interaction between `elevation` and `landuseC`. In *Model 3* (Table 1), we revert to having `forest` as the reference level and include the interaction between `elevation` and `landuseC`.

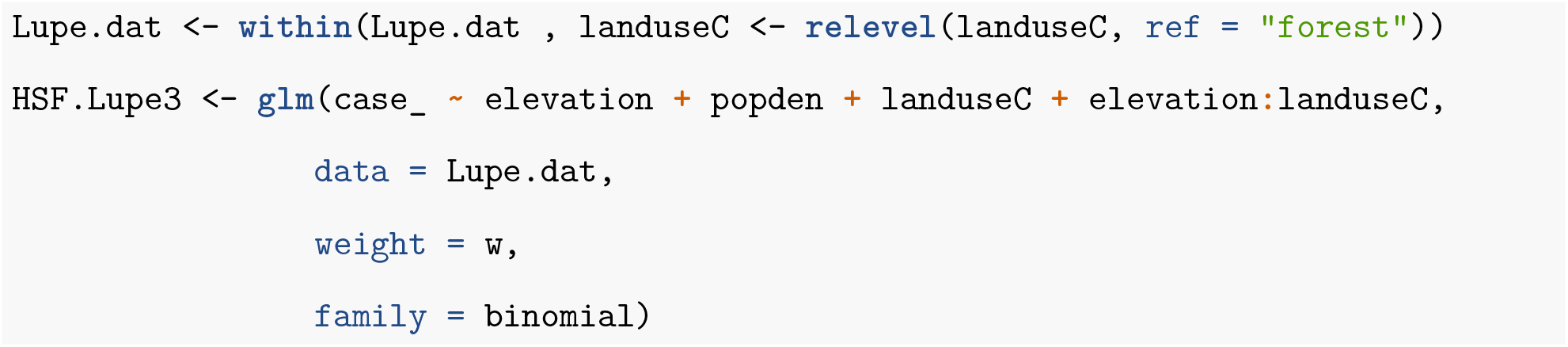

**Figure 3:**
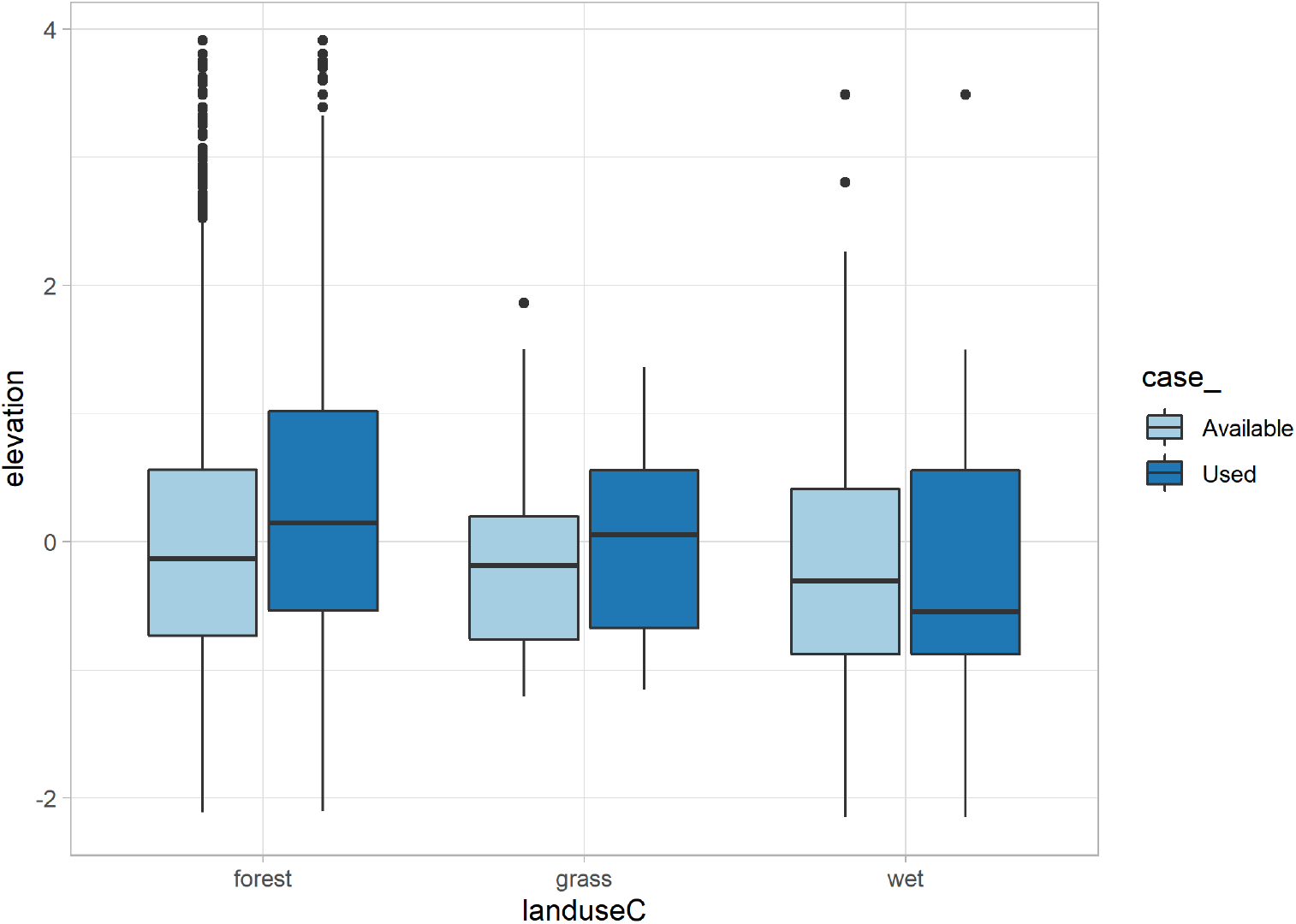
Distribution of `elevation` at used and available locations within each of 3 landcover types.

Using this syntax, R creates two new variables `elevation`:`landuseCgrass` equal to `elevation` when `landuseC` is grass and is 0 otherwise, and `elevation`:`landuseCwet` equal to `elevation` when `landuseC` is `wet` and is 0 otherwise. The coefficients associated with these predictors quantify the change in slope (i.e., change in the effect of `elevation`) when the locations fall in `grass` or `wet`, relative to the slope when the locations fall in `forest`. Starting from eqn. (6) and using the estimates for *Model 3* in Table 1, we can easily derive that the relative intensity of use of two equally available locations that differ by 1 SD unit of `elevation` is equal to exp(0.313) = 1.37 when the two locations are in `forest`, exp(0.313 + 0.112) = 1.53 when the locations are in `grass`, and exp(0.313 − 0.499) = 0.83 when the locations are in `wet` habitat. Thus, we might conclude that Lupe would select for higher `elevation`s when in `forest` or `grass`, but avoid higher elevations when in `wet`. Alternatively, we can consider how `elevation` changes Lupe’s view of the different landcover categories, noting that *β*_*grass*_ = −1.471 + 0.112`elevation` and *β*_*wet*_ = 0.183 − 0.499`elevation`. Thus, we see that Lupe’s relative avoidance of `grass` (relative to `forest`) and selection for `wet` (relative to `forest`) both decline with `elevation`, and Lupe’s inherent ranking of these 3 habitat types will change as `elevation` increases. Both interpretations are statistically correct; the analyst chooses which one to use based on the ecological motivations for the analysis (the narrative sensu Otto & Rosales, 2020).

### Non-Linear Effects and Other Considerations

When building models, it is important to consider the functional relationships between different environmental characteristics and habitat use. For example, we may classify available predictors based on whether they represent resources (higher values are generally preferable), risks (lower values are generally preferable), or conditions (values that are not too high or too low are preferable) (e.g., Matthiopoulos et al., 2015, 2020a). It is often useful to allow for non-linear effects of conditions by including quadratic terms or using a set of spline basis functions. In either case, we end up requiring multiple coefficients to capture how the intensity of use changes with the environmental predictor. Consider, for example, that we could include a quadratic term to model the effect of `elevation`, with the expectation of a unimodal habitat-selection function with respect to `elevation`. Estimating the relative use of locations *s*_1_ and *s*_2_ that differ in their values of `elevation` but are otherwise equivalent would be straightforward using eqn. (6) - we would just need to calculate the ratio of relative intensities using coefficients for `elevation` and `elevation`^2^:

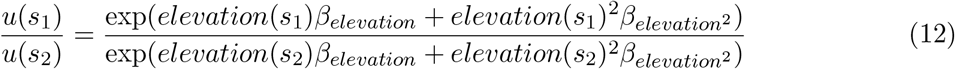

Avgar et al. (2017) provide simple formulas for calculating relative intensities under a number of different scenarios (e.g., models with quadratic polynomials, log-transformed covariates, and models with interactions). The `log_rss` function in the `amt` package (Signer et al., 2019) relies on R’s generic `predict` function to aid the user in calculating the log relative intensity for any combination of model structure and two alternative locations; its use is illustrated in Supplementary Appendix B. Understanding how these formulas are derived, however, helps build intuition and frees the user to construct estimators and estimation targets that capture relevant quantities of specific interest.

### Statistical Independence

An important assumption of the IPP model, and hence, habitat-selection functions fitted to use-availability data via logistic regression, is that any clustering of spatial locations can be explained solely by spatial covariates. Strictly speaking, this assumption will almost never be met, particularly with modern-day telemetry studies that allow several locations to be collected on the same day. Telemetry observations close in time tend to also be close in space – i.e., telemetry observations exhibit serial dependence (Fleming et al., 2014). This serial dependence is likely to manifest itself in residual spatial autocorrelation that could be modeled using a spatial random effect or a spatial predictor constructed to account for the effects of movement constraints on habitat availability (Johnson, Hooten, & Kuhn, 2013). Models with spatial random effects are, however, more complicated and difficult to fit.

Alternatively, if telemetry observations are collected at regular time intervals, then the locations may be argued to provide a representative sample of habitat use from a specific observation window (Otis & White, 1999; Fieberg, 2007). In these cases, it may be helpful to view our estimates of the parameters in our habitat-selection function, 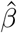, as useful summaries of habitat use for tagged individuals during these fixed time periods. Nevertheless, the assumption of independence of our locations is clearly problematic and will lead to estimates of uncertainty that are on average too small. If we are primarily interested in population-level inferences, then we may choose to ignore within-individual autocorrelation when estimating individual-specific coefficients but use a robust form of SE that treats individuals as independent when describing uncertainty in population-level parameters (e.g., using a bootstrap; Fieberg, Vitense, & Johnson, 2020) or generalized estimating equations approach (e.g., Fieberg, Rieger, Zicus, & Schildcrout, 2009; Koper & Manseau, 2009; Fieberg, Matthiopoulos, Hebblewhite, Boyce, & Frair, 2010).

## Step-Selection Functions

Step-selection functions were developed to deal with serial dependence as well as temporally varying availability distributions resulting from movement constraints (Fortin et al., 2005; Thurfjell et al., 2014). Rather than treat locations as independent and identically distributed (with availability that does not depend on time), step-selection functions model transitions, or “steps”, connecting sequential locations (Δ*t* units apart) in geographical space. The resulting redistribution kernel takes the general form:

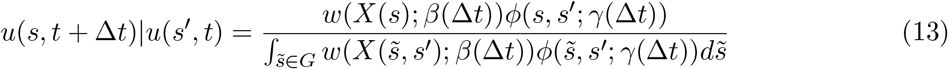

where *u*(*s, t* + Δ*t*)|*u*(*s*’, *t*) gives the conditional probability of finding the individual at location *s* at time *t* + Δ*t* given it was at location *s*’ at time *t*, *w*(*X*(*s*); *β*(Δ*t*)) is referred to as a step-selection function, and *ϕ*(*s, s*’; *γ*(Δ*t*)) is a selection-free movement kernel that describes how the animal would move in homogeneous habitat or in the absence of habitat selection (i.e., when *w*(*X*(*s*); *β*(Δ*t*)) = a constant for all *s*). Note that we represent the parameter vectors (*β* and *γ*) as functions of the step duration (Δ*t*). This notation reflects the fact that step-selection parameters are scale dependent (i.e., different Δ*t*’s will result in different estimates of *β* and *γ*; see Avgar et al., 2016 for more details). Thus, we generally require observations to be equally spaced in time (but see Munden et al., 2020), and care must be taken when comparing inference from models fitted at different temporal resolution. When animals are observed at irregular time intervals, as with many marine species, it is possible to first fit a continuous-time movement model to the location data and then use this model to provide multiply imputed data sets that are regularly spaced in time (see e.g., McClintock, 2017).

As with habitat-selection functions, it is typical to model *w*(*X*(*s*); *β*(Δ*t*)) as a log-linear function of spatial covariates and regression parameters, *w*(*X*(*s*); *β*(Δ*t*)) = exp(*X*_1_(*s*)*β*_1_ + *… X*_*k*_(*s*)*β*_*k*_). A key difference between habitat-selection functions and step-selection functions, however, is that the latter allow the available distribution to be time-dependent and equal to *a*(*s, t* + Δ*t*) = *ϕ*(*s, s*’, *γ*(Δ*t*)). Consequently, step-selection functions allow explicit consideration of temporally dynamic environmental covariates, *X*(*s*’, *t*) and *X*(*s, t* + Δ*t*) (and, possibly, environmental covariates measured along the path between these two locations). One option that often performs well and enhances interpretability is to include habitat covariates at the start of the movement step in the model for *ϕ*, and habitat covariates at the end of the movement step in the model for *w*; we provide an example in Supplementary Appendix B. This approach allows us to separately model the effect of habitat on accessibility (through the model for *ϕ*) and selection (through the model for *w*) (Matthiopoulos, 2003), and results in a more general formulation: *w*(*X*(*s, t* + Δ*t*); *β*(Δ*t*))*ϕ*(*s, s*’; *γ*(Δ*t, X*(*s*’, *t*))). We recognize, however, that there may be covariates, often measured along a movement path (e.g., crossing of a road or passing over an extremely steep slope), that also influence accessibility but that may be best included in the model for *w*. In general, we recommend including covariates in the model for *ϕ* when they are likely to influence general movement characteristics and in the model for *w* when they are likely to influence the overall attractiveness of a more limited region of geographic space.

### Models for *ϕ*(*s, s*’; *γ*(Δ*t*))

Step-selection functions build on an early idea by Arthur, Manly, McDonald, & Garner (1996) to model time-dependent availability via a circular buffer with radius *R* centered on the previous location. Rhodes, McAlpine, Lunney, & Possingham (2015) showed that this model is equivalent to assuming:

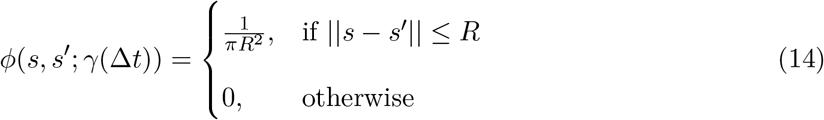

where ||*s* − *s*’|| is the Euclidean distance between locations *s* and *s*’, referred to as the *step length*. Rhodes et al. (2015) also demonstrated that circular buffers imply that individuals are more likely to move large distances than short distances since there is more area, and thus probability, associated with outer rings of the circle. Instead, they suggested using an exponential distribution to accommodate right-skewed step-length distributions and a tendency for animals to make shorter rather than longer movements:

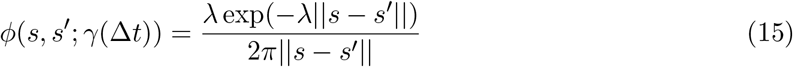

Rather than specify a model directly in terms of *ϕ*(*s, s*’; *γ*(Δ*t*)), it is more common to see movement kernels specified in terms of the distribution of step lengths, *d* = ||*s* − *s*’||, and turn angles (changes in direction from the previous bearing), *θ*. In the sections that follow, we will let *g*(*d*; *γ*_*d*_(Δ*t*)) and *f*(*θ*; *γ*_*θ*_(Δ*t*)) represent step-length and turn-angle distributions, respectively. Step-selection analyses frequently use either an exponential or gamma distribution for *g*(*d*; *γ*_*d*_(Δ*t*)). Turn angles may be assumed to be uniformly distributed as in Arthur et al. (1996) and Rhodes et al. (2015). Alternatively, circular distributions, such as the von Mises distribution or wrapped Cauchy or Weibull distributions, allow for a mode at 0 and can thus accommodate correlated movements (i.e., sequential steps are assumed, on average, to follow in the same direction as the previous step).

Although step-length and turn-angle distributions are typically assumed to be independent, animals commonly exhibit a mix of of temporally persistent movement behaviors, ranging between high-displacement movements (e.g., when traveling between habitat patches, migrating, or dispersing) and low-displacement movements (e.g., during foraging or resting bouts). If positional data are collected more frequently than the occurrence of behavioral switches, we might expect a negative cross-correlation between step lengths and turn angles (moving far is likely to coincide with moving straight) and a positive auto-correlation between the current and previous step lengths and turn angles. Moreover, as implied by the more flexible formulation, *w*(*X*(*s, t* +Δ*T*); *β*(Δ*t*))*ϕ*(*s, s*’; *γ*(Δ*t, X*(*s*’, *t*))), both step-length and turn-angle distribution may shift as a function of spatial and/or temporal covariates such as habitat permeability (e.g., terrain ruggedness, snow depth, or vegetation density), time of day, season, and predation risk (Avgar, Mosser, Brown, & Fryxell, 2013; Avgar et al., 2016). Thus, although *ϕ* is a “selection-free” movement kernel, it may still depend on environmental or temporal covariates, and hence, may vary through space and time, resulting in both auto- and cross-correlations in step attributes.

Cross-correlation between step lengths and turn angles is difficult to model with common statistical distributions, but could be accommodated using copulae (Durante & Sempi, 2010). Alternatively, one could resample (i.e., bootstrap) step length and turn angle pairs, (*d*_*t*_, *θ*_*t*_), to preserve any correlation that is present in the data (Fortin et al., 2005). Although we generally find the bootstrap appealing (Fieberg et al., 2020), it has limitations in this context. In particular, the observed distribution of step lengths and turn angles will reflect both inherent movement characteristics of the species (captured by *ϕ*) as well as habitat selection (captured by *w*). Using the observed steps as a non-parametric model for *ϕ* without adjustment for the effect of *w* can result in biased estimates of *β* (Forester, Im, & Rathouz, 2009). We will return to this point in the next section. As mentioned previously (see **Statistical Independence**), and regardless of the source of correlation, it may be preferable to calculate robust SEs by treating individuals as the relevant sampling unit when performing population-level inference (e.g., Prima, Duchesne, & Fortin, 2017). Lastly, cross- and auto-correlations in step lengths and turn angles, as well as their dependencies on various temporal or environmental characteristics, could be modeled parametrically using an integrated step-selection function (Avgar et al., 2016). To do so, we need to include appropriate statistical interactions (e.g., between concurrent and previous step lengths/turn angles and between these step-attributes and environmental or temporal covariates). We discuss this process further below, and provide examples in the Supplementary Appendix B. See also Prokopenko, Boyce, & Avgar (2017), Scrafford, Avgar, Heeres, & Boyce (2018), and Dickie et al. (2020).

### Estimation of Movement and Habitat-Selection Parameters

Although it is possible to simultaneously estimate movement (*γ*) and habitat-selection (*β*) parameters using maximum likelihood (e.g., Rhodes et al., 2015) or Bayesian methods (e.g., Johnson et al., 2008), this is rarely done in practice as it would require custom-written code. Instead, it is common to use the following approach:

1. Estimate or approximate *ϕ*(*s, s*’; *γ*(Δ*t*)) using observed step lengths and turn angles, giving 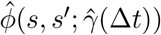.
2. Generate time-dependent available locations by simulating potential movements from the previously observed location, *u*(*t, s*’). Similar to applications of HSFs, it is up to the user to decide how many available locations to sample for each used location, and, due to similar considerations (properly approximating the availability domain: *a*(*s, t* + Δ*t*) = *ϕ*(*s, s*’; *γ*(Δ*t*)), the more points the merrier.
3. Estimate *β* using conditional logistic regression, with strata formed by combining time-dependent used and available locations.

If we knew *ϕ*(*s, s*’, *γ*(Δ*t*)) and could simulate directly from it (skipping step 1), then this approach would provide unbiased estimates of *β* (Forester et al., 2009). However, as mentioned in the previous section, estimating the selection-free movement kernel, *ϕ*(*s, s*’; *γ*(Δ*t*)), from observed steps without adjusting for habitat selection, via *w*(*X*(*s*); *β*(Δ*t*)), can lead to biased estimates of *γ* and *β*.

Forester et al. (2009) considered the case where the step-length distribution, *g*(*d, γ*_*d*_), is given by an exponential distribution with unknown parameter, *λ*. They showed that estimating *λ* directly from the observed distribution of step lengths (without adjusting for the effect of *w*(*X*(*s*); *β*(Δ*t*))), and then proceeding with steps 2 and 3 results in a biased estimators of *β*. Forester et al. (2009) also showed that the bias (if *g*(*d, γ*_*d*_) is given by an exponential distribution) is eliminated if log(*d*_*t*_) is included as a predictor in the model. Avgar et al. (2016) further showed that the coefficient associated with log(*d*_*t*_) could be used to modify 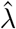, leading to an unbiased estimator of *λ* and thus, *g*(*d, γ*_*d*_). In addition, they showed how similar adjustments could be used to obtain unbiased estimators of step-length (*γ*_*d*_) and habitat-selection (*β*) parameters when the distribution of step lengths is given by a gamma, half-normal, or log-normal distribution. Similarly, Duchesne, Fortin, & Rivest (2015) showed that including cos(*θ*) as a predictor can lead to unbiased estimators of turn angle parameters (*γ*_*θ*_) when the distribution of turn angles follows a von Mises distribution. All of these adjustments are available in the `amt` package (Signer et al., 2019). Avgar et al. (2016) coined the term *integrated* step-selection analysis to emphasize that these results provide new opportunities to model both movement and habitat selection via tried and true statistical software for fitting conditional logistic regression models.

In Supplementary Appendix B, we provide a “How to” guide for implementing an integrated step-selection analysis using the `amt` package in R (R Core Team, 2019; Signer et al., 2019). Conducting an integrated step-selection analysis requires, in addition to the 3 steps outlined in this section, that we add a fourth step that re-estimates the movement parameters in *ϕ*(*s, s*’; *γ*(Δ*t*)) using regression coefficients associated with movement characteristics (e.g., log(*d*_*t*_), *cos*(*θ*)). This last step adjusts the parameters in *ϕ*(*s, s*’; *γ*(Δ*t*)) to account for the effect of habitat selection when estimating the movement kernel (Avgar et al., 2016), and is hence unnecessary if no inference about movement is being made. The details of how to carry on these adjustments are provided in Supplementary Appendix C and in Avgar et al. (2016). Importantly, interactions may be included between movement characteristics (e.g., log(*d*_*t*_), *cos*(*θ*)) and environmental covariates, *X*(*s*’, *t*), to allow the movement kernel to depend on the environment. When interactions are included, step 4 results in a movement kernel, *ϕ*(*s, s*’; *γ*(Δ*t, X*(*s*’, *t*))), that depends on the habitat the animal is in at the start of the movement step (Fig. 4).

**Figure 4:**
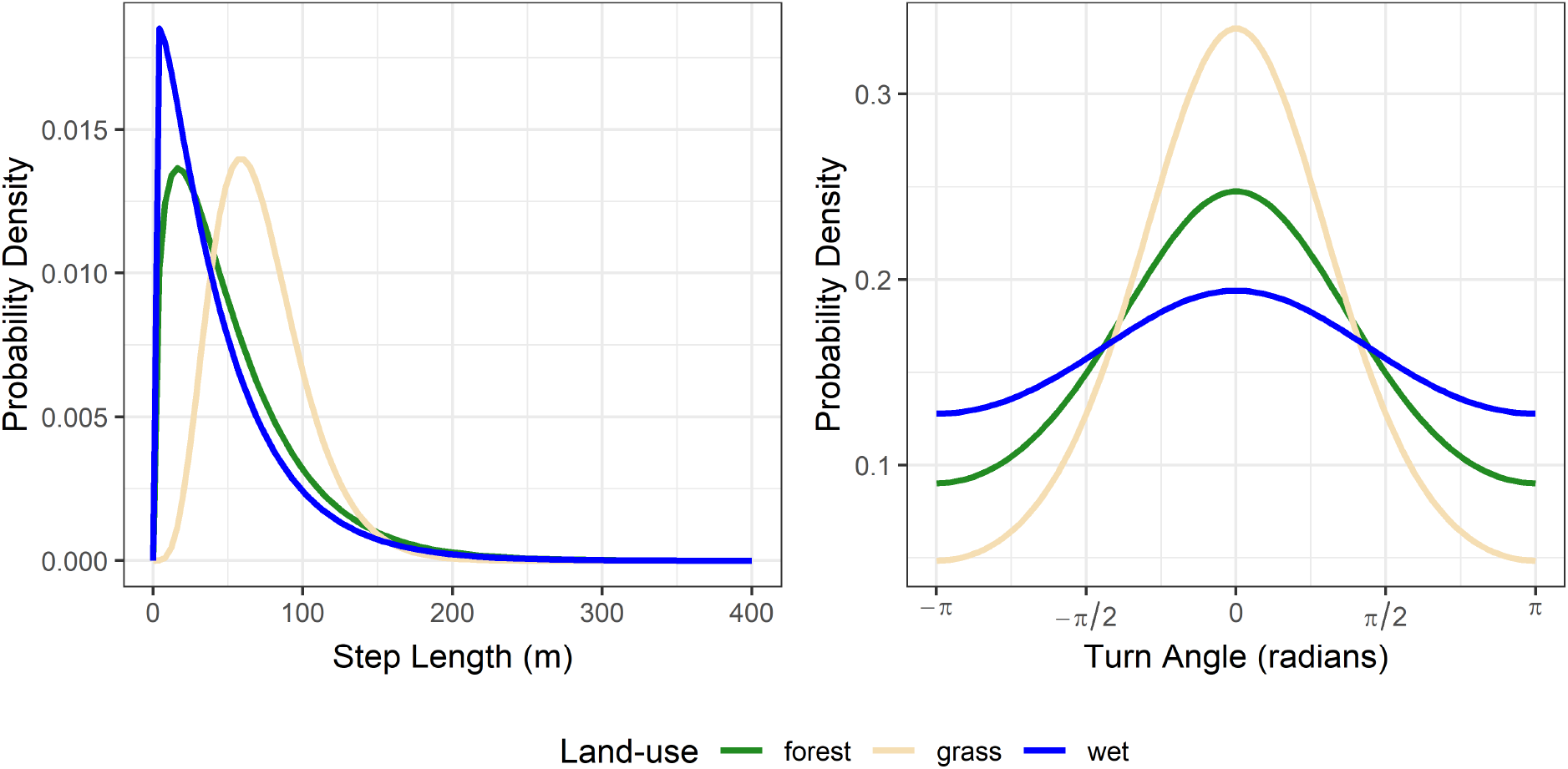
Step-length and turn-angle distributions from an integrated step-selection analysis applied to Lupe’s location data (see Supplementary Appendix B). The conditional logistic regression model included interactions between movement characteristics (step length, log step length, and cosine of the turn angle) and the landuse category Lupe was in at the start of the movement step. We see that Lupe tends to take larger, more directed steps when in `grass` and slower and more tortuous steps in `wet` habitat.

### Interpretation of Parameters in an Integrated Step-Selection Analysis

The habitat-selection parameters in an SSF can be interpreted in the same way as habitat-selection parameters in HSFs (i.e., as relative intensities, assuming locations are equally available and differing in terms of a single habitat covariate). Hence, the *ln*(*RSS*) expressions in Avgar et al. (2017), and the `log_rss` function in `amt`, are suitable for calculating and interpreting the effects of the various habitat covariates. However, it is important to recognize that the used and available distributions in step-selection analyses are dynamic and non-uniform in space. In particular, they depend an individual’s current location and movement tendencies (as well as the observed time scale determined by Δ*t*; Barnett & Moorcroft, 2008; Signer, Fieberg, & Avgar, 2017). Thus, questions that require integrating intensities over space (e.g., eqn. (10)) are more difficult to address. Possible solutions include using simulation modeling (Signer et al., 2017), solving the master equation (formed by multiplying the right hand side of eqn. (13) by *u*(*s*’, *t*) and then integrating over *G* with respect to *s*’) for its steady state (Potts et al., 2014a, 2014b), or in some cases, translating the fitted model into a partial differential equation model with analytical steady-state distribution (Potts & Schlägel, 2020). We also note that alternative modeling frameworks exist with parameters that directly describe relative intensities of use at both fine and coarse scales (e.g., Michelot et al., 2019b, 2019a; Michelot, Blackwell, Chamaillé-Jammes, & Matthiopoulos, 2020). These new analytical developments hold exciting promises to bridge the micro scale of animal movement behavior with the macro scale of animal spatial distribution but are more computationally challenging to implement. Most importantly, biologists need to be aware that parameters that describe habitat selection at local and macro scales may differ, and thus, extra steps may be required to translate movement dynamics captured by integrated step-selection analyses to the coarser scales typically modeled with traditional habitat-selection functions. The amt package has a basic capacity to simulate the utilization distribution based on a parameterized integrated step-selection function (Signer et al., 2017), and we expect this approach to become more flexible in the near future, allowing users to forecast not only steady-state utilization distributions but also transient movement patterns such as migration and dispersal.

Using an integrated step-selection approach (e.g., as in Fig. 4), it is also possible to draw ecological inference using the selection-free movement kernel. For example, the fitted step-length and turn-angle distributions can tell us how much more likely an animal is to take large versus small steps or to turn left or right relative to moving straight. We can also calculate moments of these distributions under different environmental conditions, which can be informative when our models include interactions between movement characteristics and environmental predictors. For example, we could calculate the expected selection-free displacement rates (and/or directionality) as function of local snow depth (that is, if snow depth was included in our model as an interaction with step length). Once the selection-free movement parameters are obtained, one can use them to calculate various aspects of the (theoretical) distributions of step lengths and turn angles, such as the mean, the median, or the 95% confidence bounds (see Supplementary Appendix B for examples).

## Discussion

We have highlighted how connecting habitat-selection functions to IPP models and weighted distribution theory helps with interpreting parameters in habitat-selection functions using simple examples. We have also reviewed step-selection functions and demonstrated how to estimate movement and habitat-selection parameters when conducting an integrated step-selection analysis using the `amt` package. So far, we have focused on interpreting results when analyzing data from a single individual. We end with a brief discussion addressing statistical dependencies, particularly when analyzing data from multiple individuals, along with issues related to model transferability and parameter sensitivity to changes in habitat availability and species population density.

### Statistical Dependencies

Earlier, we highlighted the importance of statistical independence as it applies to individual locations when estimating habitat-selection functions. We also noted that step-selection analyses typically assume step lengths and turn angles are independent of each other and also over time, though it is possible to account for these correlations using appropriate interactions (e.g., between step length at time *t* and time *t* − 1, step length and turn angle both at time *t*). It would be nice to have multivariate distributions available that are capable of describing correlated step lengths and turn angles and any inherent autocorrelation. It is plausible, however, that models that allow movement parameters to vary by habitat type, using interactions between step length, turn angle, and habitat covariates, will be able to account for much of the autocorrelation and cross-correlation (between step lengths and turn angles) present in the data. Similarly, autocorrelation and cross-correlations may be accommodated by models that include a (possibly latent) behavioral state, with movement and habitat-selection parameters that are state-dependent (Nicosia, Duchesne, Rivest, Fortin, & others, 2017; Suraci et al., 2019).

In addition to cross-correlation between step lengths and turn angles and serial dependencies, individuals living in different environments may exhibit different habitat-selection patterns, and thus, repeated observations on the same set of individuals will induce further statistical dependencies. A simple strategy for dealing with repeated measures when individuals can be assumed to be independent is to fit models to individual animals and then treat the resulting coefficients as data when inferring population-level patterns (Murtaugh, 2007; Fieberg et al., 2010). For example, sample means of the regression coefficients can be used to characterize average habitat-selection parameters. Estimating among-animal variability is trickier due to sampling error; naively ignoring sampling error will lead to a positive bias in estimates of among-animal variability, but more formal two-step methods can address this issue (Craiu, Duchesne, Fortin, & Baillargeon, 2011, 2016; Dickie et al., 2020). Alternatively, generalized linear mixed models with random coefficients can be used to quantify among-animal variability in habitat-selection analyses (Muff, Signer, & Fieberg, 2020).

Although it is possible to conduct integrated step-selection analyses with hierarchical models containing random effects, we have much to learn about how these approaches perform in practice. For example, Muff et al. (2020) found that parameters describing among-animal variability in habitat-selection parameters were biased low when movement characteristics were included in the model. Mixed-effect models with random coefficients are also “parameter hungry”, requiring *p*(*p* + 1)/2 variance and covariance parameters to be estimated, where *p* is the number of random coefficients. Models that allow all coefficients to be animal-specific and to covary are thus likely to be computationally challenging to fit and problematic for small data sets containing only a few individuals. For this reason, Muff et al. (2020) assumed coefficients did not covary in their applied examples. In the context of our fisher analysis, this equates to assuming that knowing an individual’s coefficient for `popden` tells us nothing about that animal’s parameters for `elevation` or `landuseC` variables. For categorical variables, it is natural to expect parameters to have a negative covariance (since, for example, spending more time in `forest` must come at the expense of spending less time in other landuse categories). Research evaluating the performance of mixed-effect step-selection analyses under various data-generating scenarios would be helpful for evaluating robustness to assumption violations (e.g., those regarding the distribution of random parameters).

### Sensitivity of Selection Coefficients to Species Population Density and Habitat Availability

Before concluding, we feel it is important to briefly discuss the oft observed pattern of density and availability dependence in habitat-selection inference (Mysterud & Ims, 1998; Matthiopoulos, Hebblewhite, Aarts, & Fieberg, 2011; Matthiopoulos et al., 2015, 2020a). Density-dependent inference may be observed when the same analysis is applied to individuals or populations of the same species, under similar environmental conditions, but at different population densities. Availability dependence (also referred to as a “functional response”) may be observed when the same analysis is applied to individuals or populations of the same species, which experience different landscape-scale resource or habitat availabilities. For example, van Beest, McLoughlin, Mysterud, & Brook (2016) found that individual elk display availability-dependent habitat-selection patterns (switching from selection to avoidance of certain habitats as function of the availability of these habitats within their home range), but that the strength of this functional response depended on elk population density. Such context dependencies are in fact so common that we do not know of a single instance where researchers were looking for them and failed to find them. Recently, Avgar, Betini, & Fryxell (2020) showed that such context dependencies in habitat-selection patterns are expected to emerge even under the simplest theoretical model of an Ideal Free Distribution (Fretwell, 1969). Thus, habitat-selection models often have poor predictive capacity when transferred across different study areas, or even within the same area over time (e.g., Torres et al., 2015). Yet, these differences may also be exploited; modeling frameworks that leverage data from multiple environments and across a range of population densities can potentially increase predictive capabilities (Matthiopoulos et al., 2019). As with any other attempt to model complex ecological data, critical evaluation of model performance for both within and out-of-sample data is essential (Fieberg, Forester, et al., 2018).

## Authors’ Contributions

JF developed the idea for the review, led the writing of the manuscript, and drafted the initial version of Supplementary Appendix A; B.S. and J.S. drafted the initial version of Supplementary Appendix B; B.S. and T.A. drafted the initial version of Supplementary Appendix C. All authors contributed critically to the manuscript text and Supplementary Files, and gave final approval for publication.

## Acknowledgements

We thank J.R. Potts for helpful comments that improved the manuscript. JF received partial salary support from the Minnesota Agricultural Experimental Station. TA received partial salary support from the Utah Agricultural Experimental Station.

## Data Availability

All of the data used in this paper are available from within the amt package (Signer et al., 2019).

### Box 1: Overview of Habitat-Selection Functions (HSFs)

- Habitat-selection functions (HSFs; historically referred to as ‘resource-selection functions’; Boyce & McDonald, 1999) provide a framework for linking locations of individual animals to important features of their environment (i.e., resources, risks, and environmental conditions).
- Exponential HSFs, the most common HSF in the literature, take the form *w*(*X*(*s*); *β*) = exp(*X*_1_(*s*)*β*_1_ + *… X*_*k*_(*s*)*β*_*k*_); where the *X*_1_(*s*)*, …, X*_*k*_(*s*) are *k* environmental predictors associated with location *s*, and the *β*_1_*, …, β*_*k*_ are parameters to be estimated.
- Parameters in HSFs are typically estimated using logistic regression, but with use-availability data rather than presence-absence data. The use of logistic regression to model use-availability data has created significant confusion in the literature.
- Inhomogeneous Poisson Point-process Models (IPPs) and Weighted Distribution Theory provide suitable frameworks for interpreting HSF parameters estimated using logistic regression. These frameworks require that users include sufficient available points to ensure parameter estimates converge to stable values (Figure 2; Warton & Shepherd, 2010). In addition, available points should be assigned large weights when fitting logistic regression models (Fithian & Hastie, 2013).
- For continuous predictors, *X*_*j*_, exponentiated HSF coefficients, exp(*β*_*j*_), quantify the relative intensity of use of locations that differ by 1 unit of *X*_*j*_, but are otherwise equivalent (i.e., they are assumed to be equally available and to have equivalent values for all other predictor variables).
- For categorical predictors, *X*_*j*_, exponentiated HSF coefficients, exp(*β*_*j*_), quantify the relative intensity of use of locations in category *j* relative to locations in a reference category, assuming both categories are equally available and that the locations do not differ with respect to other predictors.

### Box 2: Overview of Step-Selection Analyses

- Step-selection analyses model transitions or “steps” connecting sequential locations in geographical space using a selection-free movement kernel, *ϕ*, multiplied by a habitat-selection kernel, *w*. Available locations are dynamic in space and time, with availability determined by the previous location and the animal’s selection-free movement kernel.
- The selection-free movement kernel describes how the animal would move in homogeneous habitat or in the absence of habitat selection.
- Movement and habitat-selection parameters are typically estimated in a multi-step process:

1. preliminary movement parameters are estimated using observed step lengths and turn angles;
2. time-dependent availability distributions are generated by simulating potential movements from the previously observed location;
3. habitat-selection parameters are estimated using conditional logistic regression, with strata formed by combining time-dependent used and available locations;
4. if movement characteristics (e.g. log step-length, cosine of the turn angle) are included in the model, parameters associated with these characteristics can be used to update the preliminary movement parameters from step 1. Including movement characteristics in the model can reduce bias in the habitat-selection parameters (Forester et al., 2009) and improve estimates of movement parameters (Avgar et al., 2016).
- Interactions between movement characteristics (e.g. log step-length, cosine of the turn angle) and environmental covariates may be included in the conditional logistic regression model to allow the movement kernel to depend on the environment.
- Habitat-selection parameters can be interpreted in terms of relative intensities of use, assuming locations are equally available and differing in terms of a single habitat covariate. However, parameters that describe habitat-selection at local and macro scales may differ, and extra steps may be required to translate movement dynamics captured by integrated step-selection analyses to the courser scales typically modeled with HSFs (e.g., Potts et al., 2014a, 2014b; Signer et al., 2017; Potts & Schlägel, 2020).

## Supporting Information

**Supplementary Appendix A**: `AppA_HSF_examples.html`, a tutorial demonstrating how to fit and interpret parameters in habitat-selection functions.

**Supplementary Appendix B**: `AppB_SSF_examples.html`, a tutorial demonstrating how to fit and interpret parameters and output when conducting an integrated step-selection analysis.

**Supplementary Appendix C**: `AppC_iSSA_movement.html`, a description of methods used to adjust ‘tentative’ parameters in step-length and turn-angle distributions for the effects of habitat selection.

