## Supplementary Appendix A for "A ‘How-to’ Guide for Interpreting Parameters in Habitat-Selection Analyses": AppA_HSF_examples.html

### Overview

In this appendix, we will walk the user through the process of fitting and interpreting habitat-selection functions using connections between logistic regression, Inhomogeneous Poisson Point-Process (IPP) Models, and weighted distribution theory.

##### Inhomogeneous Poisson Point-Process Model

The IPP model provides a simple framework for modeling the density of points in space as a log-linear function of spatial predictors through a spatially-varying intensity function, \(\lambda(s)\):

\[\begin{equation}
\log(\lambda(s)) =\beta\_0 + X\_1(s)\beta\_1 + \ldots X\_k(s)\beta\_k
\end{equation}\]

where \(s\) is a location in geographic space, and \(X\_1(s), \ldots, X\_k(s)\) are spatial predictors associated with location \(s\). The intercept, \(\beta\_0\), determines the log-density of points (within a small homogeneous area around \(s\)) when all \(X\_i(s)\) are 0, and the slopes, \(\beta\_1, \ldots, \beta\_k\), describe the effect of spatial covariates on the log density of locations in space.

##### Weighted Distribution Theory

We can think of the *habitat-selection* function, \(w(X(s); \beta)\), as providing a set of weights that takes us from the distribution of available habitat, \(a(s)\) to the distribution of used habitat, \(u(s)\):

\[\begin{equation}
u(s) = \frac{w(X(s),\beta)a(s)}{\int\_{g \in G}w(X(g),\beta)a(g) dg} \mbox{,}
\end{equation}\]

where the denominator integrates over a geographic area, \(G\), that is assumed to be available to the animal and \(g\) is a dummy variable for integration.

### Prepare Data

We will begin by preparing our data to allow us to estimate parameters in the habitat-selection function. This will involve loading our location data and covariates, sampling random available locations from an availability domain, and associating the habitat covariates with the points.

#### Load Data

First, let’s read the fisher location and spatial covariate data. These data are available as objects in the `amt` package (`amt_fisher` and `env_covar`), and we access them easily if the package is attached. The object `amt_fisher` is already of class `track_xyt` from the package `amt`. The other three objects are all of class `RasterLayer`, which can be created by reading most standard raster files with the function `raster::raster()`. For ease, we have already loaded them, but your own data will likely need to be read in using functions from other packages.

```
fisher <- amt_fisher
landuse <- amt_fisher_covar$landuse
elevation <- amt_fisher_covar$elevation
popden <- amt_fisher_covar$popden
```

In this example, we will work with data from a single individual, “Lupe”, so let’s subset the `fisher` data and sort the locations using the timestamp of the observations.

```
dat <- fisher %>% 
  filter(name == "Lupe") %>% 
  arrange(t_)
```

The raw land-use data we loaded here has numerical codes, which we will group into different landcover types using a custom-written function, `reclass_landuse()`, below. We will apply this function when preparing the data.

```
reclass_landuse <- function(x) {
  fct_collapse(factor(x),
              forest = c("30","40","50","60", "70","80", "90","100"),
               grass = c("120", "140"),
               wet = c("160"))
}
```

Although regular sampling is not required when fitting HSFs, it is a good idea to first check the sampling rate. Your device may be programmed to collect data at regular time steps, but there will typically be some small amount of variation in the time between locations.

We can check the actual sampling rate using the `summarize_sampling_rate()` function in the `amt` package:

```
summarize_sampling_rate(dat)
```

```
## # A tibble: 1 x 9
##     min    q1 median  mean    q3   max    sd     n unit 
##   <dbl> <dbl>  <dbl> <dbl> <dbl> <dbl> <dbl> <int> <chr>
## 1 0.400  1.97   2.03  8.93  3.98 1080.  48.1  3003 min
```

We can see summary statistics for the difference between locations in minutes (see the final column for the unit).

#### Generating Available Points

We will take advantage of `amt`’s piped (`%>%`) workflow here to prepare data for the HSF. This workflow will help make our code both readable and compact. Most of the functions from `amt` have sensible defaults for each argument to get you started. You can easily see these defaults by checking the help file for that function (e.g., `?random_points`) or change them as you see fit.

In this workflow, we begin with Lupe’s location data (`dat`), generate available points (`random_points()`), and attach the environmental covariates to both the observed and available points (`extract_covariates()`). Finally, we’ll use `dplyr::mutate()` to scale and center our covariates and add a weight to the background points.

```
rsf_dat <- dat %>% 
  random_points() %>% 
  extract_covariates(landuse) %>% 
  extract_covariates(elevation) %>% 
  extract_covariates(popden) %>% 
  mutate(elevation = scale(elevation)[, 1], 
         popden = scale(popden)[, 1],
         landuseC = reclass_landuse(landuse), 
         forest = landuseC == "forest",
         weight = ifelse(case_, 1, 1e3)
  )
```

Here’s what the resulting data set looks like now. Each row is a single point with all of its associated attributes. The indicator variable, `case_`, is equal to `TRUE` for the used locations and `FALSE` for random locations.

```
print(rsf_dat, n = 3, width = Inf)
```

```
## # A tibble: 33,044 x 9
##   case_       x_       y_ landuse elevation popden landuseC forest weight
##   <lgl>    <dbl>    <dbl>   <dbl>     <dbl>  <dbl> <fct>    <lgl>   <dbl>
## 1 FALSE 1783653. 2403403.      50   -0.438   1.04  forest   TRUE     1000
## 2 FALSE 1782736. 2404285.      50    0.0985  2.44  forest   TRUE     1000
## 3 FALSE 1781872. 2401973.      50    0.884  -0.295 forest   TRUE     1000
## # ... with 33,041 more rows
```

#### Availability Domain and Number of Available Points

Let’s have a closer look at what is happening under the hood when applying the `random_points()` function. This function conveniently calculates a minimum convex polygon around Lupe’s locations. It then samples 10 random locations for every used location within this polygon. These locations are meant to capture what is assumed to be available to the animal. Other methods exist for sampling available points. For example, it is possible to sample from within an outer counter of a kernel density estimate applied to Lupe’s locations or to sample using a regular grid (see `vignette("rsf")` for more options in `amt`).

A common question is, “How many available points do I need to sample?” As discussed in the main text, the available points are used to estimate the integral in the denominator of the use-availability likelihood:

\[\begin{equation}
L(\beta; s\_1, \ldots, s\_n) = \prod\_{i=1}^{y\_s}\frac{w(X(s\_i); \beta)}{\int\_{s\in G}w(X(s); \beta) ds}
\end{equation}\]

where \(w(X(s); \beta) = \exp(\beta\_1X\_1(s) + \ldots \beta\_kX\_k(s))\) is our habitat-selection function and \(y\_G\) is the number of used points in our data set.

Below, we evaluate how \(\hat{\beta}\) changes as we increase the number of available locations from 1 available location per used location to 100 available locations per used location.

```
# Explore sensitivity of HSF coefficients to the number of available points
n.frac <- c(1, 5, 10, 50, 100)
n.pts <- ceiling(nrow(dat) * n.frac)
n.rep <- 20

res1 <- tibble(
  n.pts = rep(n.pts, n.rep), 
  frac = rep(n.frac, n.rep), 
  res = map(
    n.pts, ~
      dat %>% random_points(n = .x) %>% 
      extract_covariates(landuse) %>% 
      extract_covariates(elevation) %>% 
      extract_covariates(popden) %>% 
      mutate(landuseC = reclass_landuse(landuse), 
             elevation = scale(elevation), 
             popden = scale(popden),
             w = ifelse(case_, 1, 5000)) %>% 
      glm(case_ ~ elevation + popden + landuseC, 
          weight = w, data = ., family = binomial()) %>% 
      tidy()))

res1 %>% unnest(cols = res) %>% 
  mutate(terHSF.Lupe1 = recode(term, "(Intercept)"="Intercept",
                        popden = "human population density", 
                        landuseCgrass = "landuse == grass", 
                        landuseCother = "landuse == other", 
                        landuseCshrub = "landuse == shrub", 
                        landuseCwet = "landuse == wet", 
                        )) %>% 
  ggplot(aes(factor(frac), y = estimate)) +
  geom_boxplot() + facet_wrap(~ terHSF.Lupe1, scale  ="free") +
  geom_jitter(alpha = 0.2) + 
  labs(x = "Number of available points (multiplier of no. of used locations)", y = "Estimate") +
  theme_light()
```

We see that the intercept decreases as we increased the number of available points (as it is roughly proportional to the log difference between the numbers of used and available points), but the slope parameter estimates, on average, do not change much once we included at least 10 available points per used point. Thus, we conclude that, in this particular case, having 10 available points per used point is sufficient for interpreting the slope coefficients. Using more available points reduces Monte Carlo error, however, so we will proceed with 100 available points per used point.

Lets take a look at the proportion of used and available locations in each `landuseC` class using a data set created by combining our observed locations with 100 available points per used location.

```
#' Use the largest sample size here for the rest of the paper
Lupe.dat <- dat %>% 
  random_points(n = max(n.pts)) %>% 
  extract_covariates(landuse) %>% 
  extract_covariates(elevation) %>% 
  extract_covariates(popden) %>% 
  mutate(landuseC = reclass_landuse(landuse), 
         elevation = scale(elevation), 
         popden = scale(popden))

Lupe.dat %>% 
  group_by(case_, landuseC) %>% 
  summarize(n = n()) %>% 
  mutate(prop = n / sum(n), 
         label = paste0(round(prop * 100, 1), "%")) %>% 
  ggplot(aes(landuseC, prop, fill = case_, group=case_,label = label)) + 
  geom_col(position = position_dodge2()) +
  geom_text(size = 4, vjust = -0.25, position = position_dodge(width = 1)) +
  labs(x = "Land use class", y = "Proportion", fill = "case_")+
  scale_fill_brewer(palette = "Paired", name="case_", 
                    breaks=c("FALSE", "TRUE"), labels=c("Available", "Used")) +
  theme_light()
```

### Fitting and Interpetting Parameters in a Habitat-Selection Function (HSF)

As discussed in the main text, Fithian & Hastie (2013) showed that assigning “infinite weights” to available points ensures that the logistic regression estimators of slope coefficients converge to the those of the IPP model. Therefore, when fitting logistic regression models to use-availability data, we suggest assigning a large weight (say 1000 or more) to each available location and a weight of 1 to all observed locations (larger values can be used to verify that results are robust to this choice). Below, we use the `ifelse` function in R to assign these weights. We then fit a weighted logistic regression model with the same habitat covariates introduced in the main text, human population density (`popden`), elevation (`elevation`), and land-use category (`landuseC`). Note that we used the same model structure when evaluating the number of available points needed to accurately estimate the integral in the denominator of the likelihood function (see **Availability Domain and Number of Available Points**).

```
Lupe.dat$w <- ifelse(Lupe.dat$case_, 1, 5000)
HSF.Lupe1 <- glm(case_ ~ elevation + popden + landuseC, 
                 data = Lupe.dat, weight = w,
                 family = binomial(link = "logit"))
```

#### Interpreting Habitat-Selection Parameters in Terms of Relative Intensities or Rates of Use

Note that we will use the terms log relative intensity and log relative selective strength interchangeably. In this section, we provide R code supporting the relative intensity calculations in the main text (note, in Supplementary Appendix B, we demonstrate the use of the `log_rss` function in the `amt` package for more complicated situations). We begin by exploring the coefficients in the fitted model using the `summary()` function:

```
summary(HSF.Lupe1)
```

```
## 
## Call:
## glm(formula = case_ ~ elevation + popden + landuseC, family = binomial(link = "logit"), 
##     data = Lupe.dat, weights = w)
## 
## Deviance Residuals: 
##    Min      1Q  Median      3Q     Max  
## -0.279  -0.152  -0.136  -0.123   5.472  
## 
## Coefficients:
##               Estimate Std. Error z value Pr(>|z|)    
## (Intercept)   -13.1681     0.0195 -675.88  < 2e-16 ***
## elevation       0.3031     0.0165   18.36  < 2e-16 ***
## popden         -0.1829     0.0210   -8.70  < 2e-16 ***
## landuseCgrass  -1.4769     0.2780   -5.31  1.1e-07 ***
## landuseCwet     0.2499     0.1085    2.30    0.021 *  
## ---
## Signif. codes:  0 '***' 0.001 '**' 0.01 '*' 0.05 '.' 0.1 ' ' 1
## 
## (Dispersion parameter for binomial family taken to be 1)
## 
##     Null deviance: 84847  on 303403  degrees of freedom
## Residual deviance: 84433  on 303399  degrees of freedom
## AIC: 84443
## 
## Number of Fisher Scoring iterations: 9
```

We can exponentiate habitat-selection parameters to compare the relative intensity or rate of use of two locations that differ by 1 unit of the explanatory variable but are otherwise equivalent - i.e., they should be equally accessible and have identical values for all other explanatory variables.

##### Quantitative Predictors

We begin by considering two locations, both in the same landcover class and with the same associated population density, but differing by 1 unit in `elevation` (since we have scaled this variable, a difference of 1 implies that the two observations differ by 1 SD in the original units of elevation):

- Location s1: elevation = 3, popden = 1.5, landC = “wet”
- Location s2: elevation = 2, popden = 1.5, landC = “wet”

The relative use of location \(s\_1\) versus location \(s\_2\) is given by:

\[\begin{equation}
\frac{u(s\_1)}{u(s\_2)} = \frac{\exp(3\beta\_{elevation}+ 1.5\beta\_{pop\\_den}+0\beta\_{grass}+1\beta\_{wet})a(s\_1)}{\exp(2\beta\_{elevation}+ 1.5\beta\_{pop\\_den}+0\beta\_{grass}+1\beta\_{wet})a(s\_2)} = \exp(\beta\_{elevation})\frac{a(s\_1)}{a(s\_2)}
\end{equation}\]

Further, if we assume both locations are equally available, then \(a(s\_1)=a(s\_2)\), and this ratio simplifies to \(\exp(\beta\_{elevation}) = \frac{\lambda(s\_1)}{\lambda(s\_2)}\). Thus, this ratio is also equivalent to the ratio of intensity functions for \(s\_1\) and \(s\_2\).

```
exp(3 * coef(HSF.Lupe1)["elevation"] + 
      1.5 * coef(HSF.Lupe1)["popden"] + coef(HSF.Lupe1)["landuseCwet"]) /
  exp(2 * coef(HSF.Lupe1)["elevation"] + 1.5 * coef(HSF.Lupe1)["popden"] + 
        coef(HSF.Lupe1)["landuseCwet"])
```

```
## elevation 
##      1.35
```

```
exp(coef(HSF.Lupe1)["elevation"])
```

```
## elevation 
##      1.35
```

We get the same ratio whenever comparing any two locations that differ by 1 unit of `elevation`, provided they are equally available and have the same values for `popden` and `landuseC`.

```
exp(11 * coef(HSF.Lupe1)["elevation"] + 1.5 * coef(HSF.Lupe1)["popden"] + 
      coef(HSF.Lupe1)["landuseCwet"]) /
  exp(10 * coef(HSF.Lupe1)["elevation"] + 1.5 * coef(HSF.Lupe1)["popden"] + 
        coef(HSF.Lupe1)["landuseCwet"])
```

```
## elevation 
##      1.35
```

```
exp(111 * coef(HSF.Lupe1)["elevation"] + 1.5 * coef(HSF.Lupe1)["popden"] + 
      coef(HSF.Lupe1)["landuseCwet"]) /
  exp(110 * coef(HSF.Lupe1)["elevation"] + 1.5 * coef(HSF.Lupe1)["popden"] + 
        coef(HSF.Lupe1)["landuseCwet"])
```

```
## elevation 
##      1.35
```

```
exp(-4 * coef(HSF.Lupe1)["elevation"] + 1.5 * coef(HSF.Lupe1)["popden"] + 
      coef(HSF.Lupe1)["landuseCwet"]) /
  exp(-5 * coef(HSF.Lupe1)["elevation"] + 1.5 * coef(HSF.Lupe1)["popden"] + 
        coef(HSF.Lupe1)["landuseCwet"])
```

```
## elevation 
##      1.35
```

##### Categorical Predictors

Next, we compare two observations that differ in terms of their `landuseC` variable, but have the same values for `popden` and `elevation`. We again consider two locations, \(s\_1\) and \(s\_2\) assumed to be equally available.

- Location s1: elevation = 2, popden = 1.5, landuseC = “wet”
- Location s2: elevation = 2, popden = 1.5, landuseC = “forest”

Lupe’s relative intensity or rate of use of location 1 relative to location 2 is given by:

\[\begin{equation}
\frac{u(s\_1)}{u(s\_2)} = \frac{\exp(2\beta\_{elevation}+ 1.5\beta\_{pop\\_den}+0\beta\_{grass}+1\beta\_{wet})a(s\_1)}{\exp(2\beta\_{elevation}+ 1.5\beta\_{pop\\_den}+0\beta\_{grass}+0\beta\_{wet})a(s\_2)} = \exp(\beta\_{wet})\frac{a(s\_1)}{a(s\_2)}
\end{equation}\]

Again, if we assume the two locations are equally available, then \(a(s\_1)=a(s\_2)\), and the ratio simplifies to \(\exp(\beta\_{wet})\).

```
exp(coef(HSF.Lupe1)["landuseCwet"])
```

```
## landuseCwet 
##        1.28
```

Thus, we conclude Lupe would be 1.28 as likely to use the location in `wet` than in `forest`, assuming these two locations are equally available.

What if we change the reference level from `forest` to `wet` and refit the logistic regression model?

```
Lupe.dat <- mutate(Lupe.dat, landuseC1 = fct_relevel(landuseC, "wet"))
levels(Lupe.dat$landuseC)
```

```
## [1] "forest" "grass"  "wet"
```

```
levels(Lupe.dat$landuseC1)
```

```
## [1] "wet"    "forest" "grass"
```

```
HSF.Lupe2 <- glm(case_ ~ elevation + popden + landuseC1, 
                 data = Lupe.dat, weight = w, family = binomial(link = "logit"))
summary(HSF.Lupe2)
```

```
## 
## Call:
## glm(formula = case_ ~ elevation + popden + landuseC1, family = binomial(link = "logit"), 
##     data = Lupe.dat, weights = w)
## 
## Deviance Residuals: 
##    Min      1Q  Median      3Q     Max  
## -0.279  -0.152  -0.136  -0.123   5.472  
## 
## Coefficients:
##                 Estimate Std. Error z value Pr(>|z|)    
## (Intercept)     -12.9182     0.1070 -120.69  < 2e-16 ***
## elevation         0.3031     0.0165   18.36  < 2e-16 ***
## popden           -0.1829     0.0210   -8.70  < 2e-16 ***
## landuseC1forest  -0.2499     0.1085   -2.30    0.021 *  
## landuseC1grass   -1.7268     0.2972   -5.81  6.2e-09 ***
## ---
## Signif. codes:  0 '***' 0.001 '**' 0.01 '*' 0.05 '.' 0.1 ' ' 1
## 
## (Dispersion parameter for binomial family taken to be 1)
## 
##     Null deviance: 84847  on 303403  degrees of freedom
## Residual deviance: 84433  on 303399  degrees of freedom
## AIC: 84443
## 
## Number of Fisher Scoring iterations: 9
```

We see that the coefficients for `elevation` and `popden` do not change, but the coefficients for the landcover classes do. As discussed in the main text, the coefficients for categorical predictors reflect use:availability ratios for each level of the predictor relative to the use:availability ratio for the reference class. In this case, the coefficient for `forest` is negative, despite Lupe using `forest` more than its availability (i.e., \(u(s, s \in forest) > a(s, s \in forest)\)) and Lupe spending more than 95% of her time in the forest! The coefficient for `forest` is negative because the use:availability ratio for `forest` is less than the use:availability ratio for the reference class, `wet`.

##### Adjusting for Differences in Habitat Availability

###### Naive Adjustment Using Overall Habitat Availability

To compare Lupe’s relative use of `forest` versus `wet` habitat, we could attempt to correct for differences in availability of these landcover classes within the MCP surrounding Lupe’s locations. One simple approach is to multiply our ratio, calculated assuming equal availability, by the ratio of overall habitat availability for `wet` relative to `forest` habitats with Lupe’s MCP.

```
# Availability of forest and wet within Lupe's MCP
a.forest <- with(Lupe.dat[Lupe.dat$case_ == 0, ], sum(landuseC == "forest")) 
a.wet <- with(Lupe.dat[Lupe.dat$case_ == 0, ], sum(landuseC == "wet"))
# Multiply by the ratio of availabilities
exp(coef(HSF.Lupe1)["landuseCwet"])*a.wet/a.forest
```

```
## landuseCwet 
##      0.0302
```

```
# Or, comparing wet to forest (instead of forest to wet)
1/(exp(coef(HSF.Lupe1)["landuseCwet"])*a.wet/a.forest)
```

```
## landuseCwet 
##        33.1
```

This calculation suggests we are \((1/0.030 = 33)\) times more likely to find Lupe in `forest` than `wet` habitat. Unfortunately, with this calculation we had to naively assume that the availability distributions for `popden` and `elevation` were the same in both `wet` and `forest` cover classes. In reality, if Lupe decides to move from `forest` to `wet`, it is likely that she will experience a change in `elevation` and `popden` too (i.e., these factors will not be held constant).

###### Adjusting using Integrated Spatial Intensities

To compare Lupe’s relative use of `forest` versus `wet` habitat, while also accounting for the effects other environmental characteristics that are associated with these habitat types, we can use integrated intensities – i.e., we can integrate the spatial utilization distribution, \(u(s)\), over all `forest` and `wet` habitats:

\[\begin{equation}
\frac{u(s, s \in forest)}{u(s, s \in wet)} = \frac{\int\_{G} u(s)I(s \in forest)ds}{\int\_{G}u(s)I(s \in wet)ds}
\end{equation}\]

where \(I(s \in forest)\) and \(I(s \in wet)\) are indicator functions equal to 1 when location \(s\) is in `forest` or `wet`, respectively (and 0 otherwise). We can estimate this ratio using:

\[\begin{equation}
\frac{\hat{u}(s, s \in forest)}{\hat{u}(s, s \in wet)} = \frac{\sum\_{i=1}^{n\_a}\hat{w}(X(s\_i); \hat{\beta})I(s\_i \in forest)}{\sum\_{i=1}^{n\_a}\hat{w}(X(s\_i); \hat{\beta}))I(s\_i \in wet)}.
\end{equation}\]

where the sum is over the distribution of available locations.

```
# Filter to include only available locations, then integrate over landuse categories
available.locs <- filter(Lupe.dat, case_ == 0)
available.locs$wx <- exp(predict(HSF.Lupe1, available.locs))
with(available.locs, sum(wx[landuseC == "forest"]) / sum(wx[landuseC == "wet"]))
```

```
## [1] 33
```

```
# Now, compare to the observed ratio of Lupe's use of forest and and wet habitat 
with(Lupe.dat[Lupe.dat$case_ == 1, ], 
     sum(landuseC == "forest") / sum(landuseC == "wet"))
```

```
## [1] 33
```

We find here that the naive adjustment and integrated spatial intensities estimate are nearly identical but this will not always be the case as will be demonstrated below, by repeating these calculations using `wet` and `grass` categories.

```
# No adjustment
exp(coef(HSF.Lupe1)["landuseCwet"]) / exp(coef(HSF.Lupe1)["landuseCgrass"])
```

```
## landuseCwet 
##        5.62
```

```
# Naive adjustment
a.grass <- with(Lupe.dat[Lupe.dat$case_ == 0, ], 
                sum(landuseC == "grass")) 
(exp(coef(HSF.Lupe1)["landuseCwet"]) / 
    exp(coef(HSF.Lupe1)["landuseCgrass"])) * a.wet / a.grass
```

```
## landuseCwet 
##        6.28
```

```
# Integrated Spatial Intensities 
# Filter to include only available locations, then integrate over landuse categories
with(available.locs, sum(wx[landuseC == "wet"]) / sum(wx[landuseC == "grass"]))
```

```
## [1] 6.77
```

In this case, the estimate using integrated spatial intensities agrees best with data - the relative amount of time that Lupe was in `wet` versus `grass` habitat is shown below:

```
# Now, compare to the observed ratio in the data set
with(Lupe.dat[Lupe.dat$case_==1, ], sum(landuseC == "wet") / 
       sum(landuseC == "grass"))
```

```
## [1] 6.77
```

### Interactions

Lastly, we consider the interaction model described in the main text. We begin by plotting the distribution of elevation values at used and available locations across the different landcover classes.

```
ggplot(Lupe.dat, aes(landuseC, elevation, fill=case_))+
  geom_boxplot() +    
  scale_fill_brewer(palette = "Paired", name="case_", 
                    breaks=c("FALSE", "TRUE"), labels=c("Available", "Used"))+ 
 theme_light()
```

We then fit a model with an interaction between `elevation` and `landuseC`:

```
HSF.Lupe3 <- glm(
  case_ ~ elevation + popden + landuseC + elevation:landuseC,
  data = Lupe.dat, 
  weight=w,
  family = binomial)
summary(HSF.Lupe3)
```

```
## 
## Call:
## glm(formula = case_ ~ elevation + popden + landuseC + elevation:landuseC, 
##     family = binomial, data = Lupe.dat, weights = w)
## 
## Deviance Residuals: 
##    Min      1Q  Median      3Q     Max  
## -0.265  -0.152  -0.136  -0.123   5.492  
## 
## Coefficients:
##                         Estimate Std. Error z value Pr(>|z|)    
## (Intercept)             -13.1715     0.0196 -673.52  < 2e-16 ***
## elevation                 0.3126     0.0166   18.79  < 2e-16 ***
## popden                   -0.1862     0.0211   -8.83  < 2e-16 ***
## landuseCgrass            -1.4705     0.2779   -5.29  1.2e-07 ***
## landuseCwet               0.1832     0.1161    1.58     0.11    
## elevation:landuseCgrass   0.1115     0.3797    0.29     0.77    
## elevation:landuseCwet    -0.4985     0.1268   -3.93  8.4e-05 ***
## ---
## Signif. codes:  0 '***' 0.001 '**' 0.01 '*' 0.05 '.' 0.1 ' ' 1
## 
## (Dispersion parameter for binomial family taken to be 1)
## 
##     Null deviance: 84847  on 303403  degrees of freedom
## Residual deviance: 84417  on 303397  degrees of freedom
## AIC: 84431
## 
## Number of Fisher Scoring iterations: 9
```

We see that R created two new variables `elevation:landuseCgrass` equal to `elevation` when `landuseC` is `grass` and is 0 otherwise and `elevation:landuseCwet` equal to `elevation` when `landuseC` is `wet` and is 0 otherwise. The coefficients associated with these predictors quantify the change in slope (i.e., change in the effect of `elevation`) when the locations fall in `grass` or `wet` relative to the slope when the locations fall in `forest`. Using these results, we can easily derive that the relative intensity of use of two equally available locations that differ by 1 SD unit of `elevation` is equal to \(\exp(0.313)=1.37\) when the two locations are in `forest`, \(\exp(0.313+0.112)= 1.53\) when the locations are in `grass`, and \(\exp(0.313-0.499)= 0.83\) when the locations are in `wet` habitat. Thus, Lupe selects for higher `elevation` when in `forest` or `grass`, but avoids higher elevations when in `wet`. Alternatively, we can consider how `elevation` changes Lupe’s view of the different landcover categories, noting that \(\beta\_{grass} = -1.471 + 0.112\)`elevation` and \(\beta\_{wet} = 0.183-0.499\)`elevation`. Thus, we see that Lupe’s strength of avoidance of `grass` (relative to `forest`) and selection for `wet` (relative to `forest`) both decline with `elevation`, and Lupe’s inherent ranking of these 3 habitat types will change as `elevation` increases.

### Conclusion

We have highlighted how connecting habitat-selection functions to IPP models and weighted distribution theory helps with interpreting parameters in habitat-selection functions using simple examples involving a mix of categorical and quantitative variables.

### Session

```
sessioninfo::session_info()
```

```
## - Session info ---------------------------------------------------------------
##  setting  value                       
##  version  R version 4.0.2 (2020-06-22)
##  os       Windows 10 x64              
##  system   x86_64, mingw32             
##  ui       RTerm                       
##  language (EN)                        
##  collate  English_United States.1252  
##  ctype    English_United States.1252  
##  tz       America/Chicago             
##  date     2021-01-19                  
## 
## - Packages -------------------------------------------------------------------
##  package      * version date       lib source                       
##  amt          * 0.1.3   2020-11-04 [1] Github (jmsigner/amt@ee58ca1)
##  assertthat     0.2.1   2019-03-21 [1] CRAN (R 4.0.2)               
##  backports      1.1.7   2020-05-13 [1] CRAN (R 4.0.0)               
##  bibtex         0.4.2.2 2020-01-02 [1] CRAN (R 4.0.2)               
##  blob           1.2.1   2020-01-20 [1] CRAN (R 4.0.2)               
##  broom        * 0.7.0   2020-07-09 [1] CRAN (R 4.0.2)               
##  cellranger     1.1.0   2016-07-27 [1] CRAN (R 4.0.2)               
##  checkmate      2.0.0   2020-02-06 [1] CRAN (R 4.0.2)               
##  class          7.3-17  2020-04-26 [2] CRAN (R 4.0.2)               
##  classInt       0.4-3   2020-04-07 [1] CRAN (R 4.0.2)               
##  cli            2.0.2   2020-02-28 [1] CRAN (R 4.0.2)               
##  codetools      0.2-16  2018-12-24 [2] CRAN (R 4.0.2)               
##  colorspace     1.4-1   2019-03-18 [1] CRAN (R 4.0.2)               
##  crayon         1.3.4   2017-09-16 [1] CRAN (R 4.0.2)               
##  DBI            1.1.0   2019-12-15 [1] CRAN (R 4.0.2)               
##  dbplyr         1.4.4   2020-05-27 [1] CRAN (R 4.0.2)               
##  digest         0.6.25  2020-02-23 [1] CRAN (R 4.0.2)               
##  dplyr        * 1.0.1   2020-07-31 [1] CRAN (R 4.0.2)               
##  e1071          1.7-3   2019-11-26 [1] CRAN (R 4.0.2)               
##  ellipsis       0.3.1   2020-05-15 [1] CRAN (R 4.0.2)               
##  evaluate       0.14    2019-05-28 [1] CRAN (R 4.0.2)               
##  fansi          0.4.1   2020-01-08 [1] CRAN (R 4.0.2)               
##  farver         2.0.3   2020-01-16 [1] CRAN (R 4.0.2)               
##  forcats      * 0.5.0   2020-03-01 [1] CRAN (R 4.0.2)               
##  fs             1.5.0   2020-07-31 [1] CRAN (R 4.0.2)               
##  gbRd           0.4-11  2012-10-01 [1] CRAN (R 4.0.0)               
##  generics       0.0.2   2018-11-29 [1] CRAN (R 4.0.2)               
##  ggplot2      * 3.3.2   2020-06-19 [1] CRAN (R 4.0.2)               
##  glue           1.4.1   2020-05-13 [1] CRAN (R 4.0.2)               
##  gtable         0.3.0   2019-03-25 [1] CRAN (R 4.0.2)               
##  haven          2.3.1   2020-06-01 [1] CRAN (R 4.0.2)               
##  hms            0.5.3   2020-01-08 [1] CRAN (R 4.0.2)               
##  htmltools      0.5.0   2020-06-16 [1] CRAN (R 4.0.2)               
##  httr           1.4.2   2020-07-20 [1] CRAN (R 4.0.2)               
##  jsonlite       1.7.0   2020-06-25 [1] CRAN (R 4.0.2)               
##  KernSmooth     2.23-17 2020-04-26 [1] CRAN (R 4.0.2)               
##  knitr        * 1.30    2020-09-22 [1] CRAN (R 4.0.3)               
##  labeling       0.3     2014-08-23 [1] CRAN (R 4.0.0)               
##  lattice        0.20-41 2020-04-02 [2] CRAN (R 4.0.2)               
##  lifecycle      0.2.0   2020-03-06 [1] CRAN (R 4.0.2)               
##  lubridate      1.7.9   2020-06-08 [1] CRAN (R 4.0.2)               
##  magrittr       1.5     2014-11-22 [1] CRAN (R 4.0.2)               
##  Matrix         1.2-18  2019-11-27 [1] CRAN (R 4.0.2)               
##  modelr         0.1.8   2020-05-19 [1] CRAN (R 4.0.2)               
##  munsell        0.5.0   2018-06-12 [1] CRAN (R 4.0.2)               
##  pillar         1.4.6   2020-07-10 [1] CRAN (R 4.0.2)               
##  pkgconfig      2.0.3   2019-09-22 [1] CRAN (R 4.0.2)               
##  purrr        * 0.3.4   2020-04-17 [1] CRAN (R 4.0.2)               
##  R6             2.4.1   2019-11-12 [1] CRAN (R 4.0.2)               
##  raster         3.3-13  2020-07-17 [1] CRAN (R 4.0.2)               
##  RColorBrewer   1.1-2   2014-12-07 [1] CRAN (R 4.0.0)               
##  Rcpp           1.0.5   2020-07-06 [1] CRAN (R 4.0.2)               
##  Rdpack         1.0.0   2020-07-01 [1] CRAN (R 4.0.2)               
##  readr        * 1.3.1   2018-12-21 [1] CRAN (R 4.0.2)               
##  readxl         1.3.1   2019-03-13 [1] CRAN (R 4.0.2)               
##  reprex         0.3.0   2019-05-16 [1] CRAN (R 4.0.2)               
##  rlang          0.4.7   2020-07-09 [1] CRAN (R 4.0.2)               
##  rmarkdown      2.3     2020-06-18 [1] CRAN (R 4.0.2)               
##  rstudioapi     0.11    2020-02-07 [1] CRAN (R 4.0.2)               
##  rvest          0.3.6   2020-07-25 [1] CRAN (R 4.0.2)               
##  scales         1.1.1   2020-05-11 [1] CRAN (R 4.0.2)               
##  sessioninfo    1.1.1   2018-11-05 [1] CRAN (R 4.0.2)               
##  sf             0.9-6   2020-09-13 [1] CRAN (R 4.0.2)               
##  sp             1.4-2   2020-05-20 [1] CRAN (R 4.0.2)               
##  stringi        1.4.6   2020-02-17 [1] CRAN (R 4.0.0)               
##  stringr      * 1.4.0   2019-02-10 [1] CRAN (R 4.0.2)               
##  survival       3.1-12  2020-04-10 [2] CRAN (R 4.0.2)               
##  tibble       * 3.0.3   2020-07-10 [1] CRAN (R 4.0.2)               
##  tidyr        * 1.1.1   2020-07-31 [1] CRAN (R 4.0.2)               
##  tidyselect     1.1.0   2020-05-11 [1] CRAN (R 4.0.2)               
##  tidyverse    * 1.3.0   2019-11-21 [1] CRAN (R 4.0.2)               
##  units          0.6-7   2020-06-13 [1] CRAN (R 4.0.2)               
##  utf8           1.1.4   2018-05-24 [1] CRAN (R 4.0.2)               
##  vctrs          0.3.2   2020-07-15 [1] CRAN (R 4.0.2)               
##  withr          2.2.0   2020-04-20 [1] CRAN (R 4.0.2)               
##  xfun           0.16    2020-07-24 [1] CRAN (R 4.0.2)               
##  xml2           1.3.2   2020-04-23 [1] CRAN (R 4.0.2)               
##  yaml           2.2.1   2020-02-01 [1] CRAN (R 4.0.2)               
## 
## [1] C:/Users/jfieberg/Documents/R/win-library/4.0
## [2] C:/Program Files/R/R-4.0.2/library
```
