## Supplementary Appendix B for "A ‘How-to’ Guide for Interpreting Parameters in Habitat-Selection Analyses": AppB_SSF_examples.html

### Overview

In this appendix, we will walk the user through the process of fitting and interpreting parameters and output when conducting an *integrated step-selection analysis*. Step-selection functions model transitions or “steps” connecting sequential locations (\(\Delta t\) units apart) in geographical space:

\[\begin{equation}
u(s,t+\Delta t) | u(s', t) = \frac{w(X(s); \beta(\Delta t))\phi(s, s'; \gamma(\Delta t))}{\int\_{\tilde{s}\in G}w(X(\tilde{s}, s'); \beta(\Delta t))\phi(\tilde{s},s'; \gamma(\Delta t))d\tilde{s}}
\end{equation}\]

where \(u(s, t+\Delta t) | u(s', t)\) gives the conditional probability of finding the individual at location \(s\) at time \(t+\Delta t\) given it was at location \(s'\) at time \(t\), \(w(X(s); \beta(\Delta t))\) is referred to as a step-selection function, and \(\phi(s,s'; \gamma(\Delta t))\) is a selection-independent movement kernel that describes how the animal would move in homogeneous habitat or in the absence of habitat selection (i.e., when \(w(X(s); \beta(\Delta t))=\) a constant for all \(s\)).

Integrated step-selection analyses are implemented using the following steps:

1. Estimate a tentative selection-free movement kernel, \(\phi(s,s'; \gamma(\Delta t))\), using observed step-lengths and turn angles, giving \(\hat{\phi}(s,s'; \hat{\gamma}(\Delta t))\).
2. Generate time-dependent available locations by simulating potential movements from the previously observed location: : \(a(s, t+\Delta t) = \phi(s,s'; \gamma(\Delta t))\).
3. Estimate \(\beta\) using conditional logistic regression, with strata formed by combining time-dependent used and available locations.
4. Re-estimate the movement parameters in \(\phi(s,s'; \gamma(\Delta t))\) using regression coefficients associated with movement characteristics (e.g., step-length, log step-length, and cosine of the turn angle). This step adjusts the parameters in \(\phi(s,s'; \gamma(\Delta t))\) to account for the effect of habitat selection when estimating the movement kernel (Avgar, Potts, Lewis, & Boyce, 2016).

The first two steps are accomplished in the **Format and Generate Random Steps** and **Tentative Movement Parameters** sections. The third step is accomplished in the **Fit iSSF** section. The last step is accomplished in the **Interpreting Movement Parameters** section.

### Prepare Data

We will begin by preparing our data to allow us to estimate parameters in the movement kernel, \(\phi(s,s';\gamma(\Delta t))\) and the habitat-selection function, \(w(X(s); \beta(\Delta t))\). This step will involve loading our location data and covariates, translating the locations into a series of regular steps (transitions between sequential locations spaced \(\Delta t\) units apart), sampling random available steps from a tentative statistical distribution formulated in terms of step-length distribution and turn-angle distribution, and associating the habitat covariates with the steps.

#### Load Data

First, let’s read the fisher location and spatial covariate data. These data are available as objects in the `amt` package (`amt_fisher` and `env_covar`), and we access them easily if the package is attached. The object `amt_fisher` is already of class `track_xyt` from the package `amt`, which we have already seen in the RSF examples (see Supplementary Appendix A). The other three objects are all of class `RasterLayer`, which can be created by reading most standard raster files with the function `raster::raster()`. For ease, we have already loaded them, but your own data will likely need to be read in using functions from other packages.

```
fisher <- amt_fisher
landuse <- amt_fisher_covar$landuse
elevation <- amt_fisher_covar$elevation
popden <- amt_fisher_covar$popden
```

In this example, we will work with data from a single individual, “Lupe”, so let’s subset the `fisher` data and sort the locations using the timestamp of the observations, `t_`.

```
dat <- fisher %>% 
  filter(name == "Lupe") %>% 
  arrange(t_)
```

The raw land-use data we loaded here has numerical codes, which we will group into different landcover types using a custom-written function, `reclass_landuse()`, below. We will apply this function when preparing the data.

```
reclass_landuse <- function(x) {
  fct_collapse(factor(x),
               forest = c("30","40","50","60", "70","80", "90","100"),
               grass = c("120", "140"),
               wet = c("160"))
}
```

Most step-selection analyses assume data are collected at regular time steps (for an exception, see Munden et al., 2020). Although your device may be programmed to collect data at regular time steps, there will typically be some small amount of variation in the time between locations. If we consider that variability to be negligible, we can round it off by resampling the track.

Check the actual sampling rate:

```
summarize_sampling_rate(dat)
```

```
## # A tibble: 1 x 9
##     min    q1 median  mean    q3   max    sd     n unit 
##   <dbl> <dbl>  <dbl> <dbl> <dbl> <dbl> <dbl> <int> <chr>
## 1 0.400  1.97   2.03  8.93  3.98 1080.  48.1  3003 min
```

We can see summary statistics for the difference between locations in minutes (see the final column for the unit). These tags were programmed to take a location every two minutes, but there is some variation in the time between locations, and missed locations lead to some gaps in the data. We will therefore need to resample the track when we format the data for this analysis.

#### Format and Generate Random Steps

We will take advantage of `amt`’s piped (`%>%`) workflow here to prepare data for model fitting. This workflow will help make our code both readable and compact. Most of the functions from `amt` have sensible defaults for each argument to get you started. You can easily see these defaults by checking the help file for that function (e.g., `?random_steps`) or change them as you see fit.

In this workflow, we begin with Lupe’s location data (`dat`), we resample the track to get regular time steps (`track_resample()`), convert from points to steps (`steps_by_burst()`), generate random available steps (`random_steps()`), and attach the environmental covariates to both the observed and available steps (`extract_covariates()`). Finally, we’ll use `dplyr::mutate()` to scale and center our covariates and add the additional movement covariates that we will need to conduct an integrated step-selection analysis.

```
ssf_dat <- dat %>% 
  track_resample(rate = minutes(2), tolerance = seconds(20)) %>% 
  steps_by_burst() %>% 
  random_steps() %>% 
  extract_covariates(landuse, where = "both") %>% 
  extract_covariates(elevation, where = "both") %>% 
  extract_covariates(popden, where = "both") %>% 
  mutate(elevation_start = scale(elevation_start), 
         elevation_end = scale(elevation_end),
         popden_start = scale(popden_start),
         popden_end = scale(popden_end),
         landuseC_start = reclass_landuse(landuse_start), 
         landuseC_end = reclass_landuse(landuse_end), 
         forest_start = landuseC_start == "forest",
         forest_end = landuseC_end == "forest",
         cos_ta_ = cos(ta_), 
         log_sl_ = log(sl_)
  ) %>% 
  filter(!is.na(ta_))
```

Before we proceed, we make a few notes about the code above:

- by default the `random_steps()` function fits a tentative gamma distribution to the observed step lengths and a tentative von Mises distribution to the observed turn angles. It then generates random available points by sampling step-lengths and turn angles from these fitted distributions and combining these random steps with the starting locations associated with each observed movement step.
- by including the argument `where = "both"` to the `extract_covariates()` function, we are able to extract covariates associated with both the starting and ending locations of each movement step. It is often sensible to use covariates associated with the starting location to model the effect of habitat on the movement kernel, \(\phi(s,s';\gamma(\Delta t)\), and covariates associated with the ending location to model the habitat-selection function, \(w(X(s); \beta(\Delta t))\) (Signer, Fieberg, & Avgar, 2019).

Here’s what the resulting data set looks like now. Each row is a single step with all of its associated attributes:

- `burst_` is a unique identifier associated with a set of consecutive steps that are spaced \(\Delta t\) units apart.
- `x1_` and `y1_` are the coordinates associated with the starting location of the step.
- `x2_` and `y2_` are the coordinates associated with the ending location of the step.
- `sl_` and `ta_` are the step-length and turn angle associated with the step.
- `t1_` and `t2_` are the timestamps associated with the start and end of the step.
- `dt_` is the time duration associated with the step.
- `step_id_` is a unique identifier associated with each step’s observed and random locations.
- `case_` is an indicator variable equal to `TRUE` for observed steps and `FALSE` for random steps.
- (`landuse_start`, `elevation_start`, `popden_start`) and (`landuse_end`, `elevation_end`, `popden_end`) are covariate values for (landuse categories, elevation, and population density) associated with the start and end of the movement step, respectively.
- `cos_ta_` and `log_sl` capture the cosine of the turn angle and the log of the step-length. These covariates will allow us to adjust/refine the parameters of our tentative step-length and turn-angle distributions after fitting our integrated step-selection model.

```
print(ssf_dat, n = 3, width = Inf)
```

```
## # A tibble: 16,175 x 25
##   burst_      x1_      x2_      y1_      y2_   sl_ direction_p    ta_
## *  <dbl>    <dbl>    <dbl>    <dbl>    <dbl> <dbl>       <dbl>  <dbl>
## 1      6 1783660. 1783673. 2402636. 2402664. 31.2         2.90 -1.75 
## 2      6 1783660. 1783659. 2402636. 2402633.  3.36        2.90  1.48 
## 3      6 1783660. 1783654. 2402636. 2402631.  6.92        2.90  0.907
##   t1_                 t2_                 dt_       step_id_ case_ landuse_start
## * <dttm>              <dttm>              <drtn>       <int> <lgl>         <dbl>
## 1 2010-12-17 09:05:14 2010-12-17 09:07:04 1.83 mins        1 FALSE            50
## 2 2010-12-17 09:05:14 2010-12-17 09:07:04 1.83 mins        1 FALSE            50
## 3 2010-12-17 09:05:14 2010-12-17 09:07:04 1.83 mins        1 FALSE            50
##   landuse_end elevation_start[,1] elevation_end[,1] popden_start[,1]
## *       <dbl>               <dbl>             <dbl>            <dbl>
## 1          50              -0.275            -0.810           -0.754
## 2          50              -0.275            -0.230           -0.754
## 3          50              -0.275            -0.230           -0.754
##   popden_end[,1] landuseC_start landuseC_end forest_start forest_end cos_ta_
## *          <dbl> <fct>          <fct>        <lgl>        <lgl>        <dbl>
## 1         -0.763 forest         forest       TRUE         TRUE       -0.181 
## 2         -0.763 forest         forest       TRUE         TRUE        0.0862
## 3         -0.763 forest         forest       TRUE         TRUE        0.616 
##   log_sl_
## *   <dbl>
## 1    3.44
## 2    1.21
## 3    1.93
## # ... with 16,172 more rows
```

#### Tentative Movement Parameters

Let’s have a closer look at what is happening under the hood when applying the `random_steps()` function. This function conveniently fits tentative step-length and turn-angle distributions and then samples from these distributions to generate random steps. The default arguments lead to `random_steps()` fitting a gamma distribution for the step lengths and a von Mises distribution for the turn angles.

The tentative parameters in these statistical distributions (gamma, von Mises) are stored as attributes of the resulting object. We can view the parameters of the tentative step-length distribution using:

```
sl_distr(ssf_dat)
```

```
## $name
## [1] "gamma"
## 
## $params
## $params$shape
## [1] 1.42
## 
## $params$scale
## [1] 35.9
## 
## 
## attr(,"class")
## [1] "gamma_distr" "sl_distr"    "amt_distr"   "list"
```

Similarly, we can access the parameters of the tentative turn-angle distribution using:

```
ta_distr(ssf_dat)
```

```
## $name
## [1] "vonmises"
## 
## $params
## $params$kappa
## [1] 0.484
## 
## $params$mu
## [1] 0
## 
## 
## attr(,"class")
## [1] "vonmises_distr" "ta_distr"       "amt_distr"      "list"
```

After we fit the iSSF, we will see how we can update these distributions to adjust for the influence of habitat selection when estimating parameters of the movement kernel.

### Basic iSSF

In this section, we will fit a basic iSSF, demonstrating how to interpret habitat effects and movement parameters. In the next section, we will fit more complex models and show how those models differ from the basic approach presented here.

#### Fit iSSF

We will fit a model with the same habitat covariates introduced in the main text. Suppose we hypothesize that habitat selection at the step scale depends on human population density (`popden`), elevation (`elevation`), and land-use category (`landuseC`). Furthermore, suppose we hypothesize that, in the absence of habitat selection, step lengths and turn angles follow a constant distribution, i.e., the selection-free movement kernel does not depend on covariates. We can fit that model as follows:

```
m1 <- ssf_dat %>% 
  fit_issf(case_ ~ popden_end + elevation_end + landuseC_end + 
             sl_ + log_sl_ + cos_ta_ + 
             strata(step_id_), model = TRUE)
```

Notice that our model formula includes habitat covariates (`popden`, `elevation`, and `landuseC`) and movement covariates (`sl_`, `log_sl_`, and `cos_ta_`), as well as an indicator for the column that indicates the stratum, i.e., the grouping that matches each observed step to its corresponding available steps (`step_id_`). The extra argument, `model = TRUE`, is passed along to `survival::clogit()` to tell it to store the original model data in the resulting object (`m1`). We will need this extra information later when we use our fitted model to generate predictions.

#### Interpreting Habitat-Selection Parameters

We begin by exploring the coefficients in the fitted model using the `summary()` function:

```
summary(m1)
```

```
## Call:
## coxph(formula = Surv(rep(1, 16175L), case_) ~ popden_end + elevation_end + 
##     landuseC_end + sl_ + log_sl_ + cos_ta_ + strata(step_id_), 
##     data = data, model = TRUE, method = "exact")
## 
##   n= 16175, number of events= 1155 
## 
##                        coef exp(coef)  se(coef)     z Pr(>|z|)   
## popden_end        -0.107806  0.897802  0.106930 -1.01   0.3134   
## elevation_end      0.169540  1.184760  0.059918  2.83   0.0047 **
## landuseC_endgrass -1.195003  0.302703  0.510012 -2.34   0.0191 * 
## landuseC_endwet   -0.047474  0.953635  0.369706 -0.13   0.8978   
## sl_                0.000784  1.000784  0.001306  0.60   0.5483   
## log_sl_            0.023037  1.023305  0.057365  0.40   0.6880   
## cos_ta_            0.008637  1.008675  0.045452  0.19   0.8493   
## ---
## Signif. codes:  0 '***' 0.001 '**' 0.01 '*' 0.05 '.' 0.1 ' ' 1
## 
##                   exp(coef) exp(-coef) lower .95 upper .95
## popden_end            0.898      1.114     0.728     1.107
## elevation_end         1.185      0.844     1.053     1.332
## landuseC_endgrass     0.303      3.304     0.111     0.823
## landuseC_endwet       0.954      1.049     0.462     1.968
## sl_                   1.001      0.999     0.998     1.003
## log_sl_               1.023      0.977     0.914     1.145
## cos_ta_               1.009      0.991     0.923     1.103
## 
## Concordance= 0.522  (se = 0.01 )
## Likelihood ratio test= 18.3  on 7 df,   p=0.01
## Wald test            = 17.8  on 7 df,   p=0.01
## Score (logrank) test = 18  on 7 df,   p=0.01
```

For now, we will just focus on the habitat-selection coefficients, i.e., `popden`, `elevation`, and the two `landuseC` coefficients. The interpretation of the parameters is very similar to the RSF interpretation (Supplementary Appendix A) but at a local scale with habitat availability determined by the selection-free movement kernel. We can exponentiate habitat-selection parameters in a fitted step-selection model to compare the relative rates of use of two locations that differ by 1 unit of the explanatory variable but are otherwise equivalent - i.e., they should be equally accessible and have identical values for all other explanatory variables.

##### Calculating Relative Selection Strength (RSS) for Two Locations

Calculating the relative use of location \(s\_1\) versus location \(s\_2\) is fairly straightforward when \(s\_1\) and \(s\_2\) share the same values for all but one covariate (see e.g., Supplementary Appendix A). For more complex scenarios, we have implemented a function in `amt` that will calculate the log-relative intensity (referred to as the log-Relative Selection Strength, or log-RSS, by Avgar, Lele, Keim, & Boyce, 2017) by leveraging the generic `predict()` function available for many ordinary model classes (e.g., `lm()` or `glm()`) models in `R`. Regardless of whether there are interactions or polynomial terms, the function will correctly calculate log-RSS for you. If you prefer to quantify relative-use (i.e., RSS), you can simply exponentiate the results.

Here we demonstrate the use of`log_rss()` with the simple example presented in equation 8 of the main text. Suppose we have two locations with the following covariates:

- \(s\_1\): `elevation_end = 3`, `popden_end = 1.5`, `landuseC_end = "wet"`
- \(s\_2\): `elevation_end = 2`, `popden_end = 1.5`, `landuseC_end = "wet"`

We want to calculate the log-RSS for a step ending in \(s\_1\) versus a step ending in \(s\_2\), assuming \(s\_1\) and \(s\_2\) are equally accessible to the animal.

The function `log_rss()` expects the two locations to be formatted as separate `data.frame` objects. Each `data.frame` must include all covariates used to fit the model. Furthermore, `factor` variables should have the same `levels` as the original data.

```
# data.frame for s1
s1 <- data.frame(
  elevation_end = 3,
  popden_end = 1.5,
  landuseC_end = factor("wet", levels = levels(ssf_dat$landuseC_end)),
  sl_ = 100,
  log_sl_ = log(100),
  cos_ta_ = 1)

# data.frame for s2
s2 <- data.frame(
  elevation_end = 2,
  popden_end = 1.5,
  landuseC_end = factor("wet", levels = levels(ssf_dat$landuseC_end)),
  sl_ = 100,
  log_sl_ = log(100),
  cos_ta_ = 1)
```

Now that we have specified each location as a `data.frame`, we can pass them along to `log_rss()` for the calculation. The function will return an object of class `log_rss`, which is also more generally a `list`. The `list` element `"df"` contains a `data.frame` which contains the log-RSS calculation and could easily be used to make a plot when considering relative selection strength across a range of environmental characteristics.

```
lr1 <- log_rss(m1, x1 = s1, x2 = s2)
lr1$df
```

```
##   elevation_end_x1 popden_end_x1 landuseC_end_x1 sl_x1 log_sl_x1 cos_ta_x1
## 1                3           1.5             wet   100      4.61         1
##   log_rss
## 1    0.17
```

As we will see below, this function is designed to be able to consider several locations as `x1`, relative to a single location in `x2`. Because the `[["df"]]` element of the `log_rss` object is designed for plotting, it includes the value of all the covariates at location `x1` by appending `_x1` to the end of the predictor names. The `log_rss` object also contains the original `x1` and `x2` as elements, along with the model formula.

We can see that the log-RSS for this example is 0.17, which is equivalent to \(\beta\_{elevation}\) since we have considered the simple case in which \(s\_1\) and \(s\_2\) differ by only a single unit of elevation. Now that we know how the function works, we can consider more complex cases.

Suppose we are interested in making a figure showing the relative selection strength our fisher using a range of elevations. The function `log_rss()` can accept a `data.frame` with many rows as as its first argument, `x1`, to go with a single comparison location, `x2` (i.e., `x2` must only have 1 row).

```
# Make a new data.frame for s1
s1 <- data.frame(
  elevation_end = seq(from = -2, to = 2, length.out = 200),
  popden_end = 1.5,
  landuseC_end = factor("wet", levels = levels(ssf_dat$landuseC_end)),
  sl_ = 100,
  log_sl_ = log(100),
  cos_ta_ = 1)

# data.frame for s2
s2 <- data.frame(
  elevation_end = 0, # mean of elev, since we scaled and centered
  popden_end = 1.5,
  landuseC_end = factor("wet", levels = levels(ssf_dat$landuseC_end)),
  sl_ = 100,
  log_sl_ = log(100),
  cos_ta_ = 1)

# Calculate log-RSS
lr2 <- log_rss(m1, s1, s2)

# Plot using ggplot2
ggplot(lr2$df, aes(x = elevation_end_x1, y = log_rss)) +
  geom_line(size = 1) +
  geom_hline(yintercept = 0, linetype = "dashed", color = "gray30") +
  xlab("Elevation (SD)") +
  ylab("log-RSS vs Mean Elevation") +
  theme_bw()
```

The line depicts the log-RSS between each value of \(s\_1\) relative to \(s\_2\). Note that these locations differ only in their values of `elevation`, and the log-RSS is equal to 0 when the two locations are identical (i.e., when `elevation` = 0). If you prefer to plot RSS to log-RSS, simply exponentiate the y-axis (and move your horizontal line for no selection to \(exp(0) = 1\)).

```
# Plot using ggplot2
ggplot(lr2$df, aes(x = elevation_end_x1, y = exp(log_rss))) +
  geom_line(size = 1) +
  geom_hline(yintercept = exp(0), linetype = "dashed", color = "gray30") +
  xlab("Elevation (SD)") +
  ylab("RSS vs Mean Elevation") +
  theme_bw()
```

##### Bootstrap Confidence Intervals

The RSS plots above do not give any indication of model uncertainty. One method for constructing confidence intervals associated with the RSS predictions is via non-parametric bootstrapping (Fieberg, Vitense, & Johnson, 2020). When using the argument `ci="boot"` to the `log_rss()` function, the function repeatedly resamples strata with replacement, refits the model, and calculates log-RSS. Quantiles of the bootstrap log-RSS statistics are used to construct, by default, a 95% confidence interval around the original log-RSS calculation. The size of the confidence interval and number of bootstrap iterations can be controlled by the user by changing the `ci_level` and `n_boot` arguments. Warning: this code can take some time to execute.

```
# Calculate log-RSS
lr2_ci_boot <- log_rss(m1, s1, s2, ci = "boot", ci_level = 0.95, n_boot = 1000)
```

```
## Generating bootstrapped confidence intervals...
##    Expect this to take some time... 
##  ... finished bootstrapping.
```

```
# Check the header of the data.frame
head(lr2_ci_boot$df)
```

```
##   elevation_end_x1 popden_end_x1 landuseC_end_x1 sl_x1 log_sl_x1 cos_ta_x1
## 1            -2.00           1.5             wet   100      4.61         1
## 2            -1.98           1.5             wet   100      4.61         1
## 3            -1.96           1.5             wet   100      4.61         1
## 4            -1.94           1.5             wet   100      4.61         1
## 5            -1.92           1.5             wet   100      4.61         1
## 6            -1.90           1.5             wet   100      4.61         1
##   log_rss    lwr     upr
## 1  -0.339 -0.578 -0.0755
## 2  -0.336 -0.572 -0.0747
## 3  -0.332 -0.566 -0.0739
## 4  -0.329 -0.560 -0.0732
## 5  -0.325 -0.555 -0.0724
## 6  -0.322 -0.549 -0.0717
```

You can see that the resulting `lr2_ci_boot` data.frame now has columns `lwr` and `upr`, indicating the lower and upper bounds of the confidence interval. We can now add this confidence envelope to our plot.

```
ggplot(lr2_ci_boot$df, aes(x = elevation_end_x1, y = log_rss)) +
  geom_ribbon(aes(ymin = lwr, ymax = upr), 
              linetype = "dashed", 
              color = "black", fill = "gray80", alpha = 0.5) +
  geom_line(size = 1) +
  geom_hline(yintercept = 0, linetype = "dashed", color = "gray30") +
  xlab("Elevation (SD)") +
  ylab("log-RSS vs Mean Elevation") +
  theme_bw()
```

##### Large-Sample Confidence Intervals

To avoid the computational time required to calculate bootstrap confidence intervals, you may decide that using a large-sample-based confidence interval is appropriate. We can use the standard errors from our fitted model to estimate these confidence intervals based on a normal approximation to the sampling distribution of \(\hat{\beta}\).

```
lr2_ci_se <- log_rss(m1, s1, s2, ci = "se", ci_level = 0.95)

# Check the header of the data.frame
head(lr2_ci_se$df)
```

```
##   elevation_end_x1 popden_end_x1 landuseC_end_x1 sl_x1 log_sl_x1 cos_ta_x1
## 1            -2.00           1.5             wet   100      4.61         1
## 2            -1.98           1.5             wet   100      4.61         1
## 3            -1.96           1.5             wet   100      4.61         1
## 4            -1.94           1.5             wet   100      4.61         1
## 5            -1.92           1.5             wet   100      4.61         1
## 6            -1.90           1.5             wet   100      4.61         1
##   log_rss    lwr    upr
## 1  -0.339 -0.574 -0.104
## 2  -0.336 -0.568 -0.103
## 3  -0.332 -0.562 -0.102
## 4  -0.329 -0.557 -0.101
## 5  -0.325 -0.551 -0.100
## 6  -0.322 -0.545 -0.099
```

We see, in this case, that the two confidence intervals (bootstrap, normal-based) lead to similar conclusions.

```
# Add "type" label to both predictions data.frames
lr2_ci_boot$df$type <- "Bootstrap"
lr2_ci_se$df$type <- "SE"

# Combine two prediction data.frames
lr2_ci_both <- rbind(lr2_ci_boot$df, lr2_ci_se$df)

# Plot
ggplot(lr2_ci_both, aes(x = elevation_end_x1, y = log_rss, 
                        group = factor(type), 
                        fill = factor(type))) +
  geom_ribbon(aes(ymin = lwr, ymax = upr), 
              linetype = "dashed", color = "black", alpha = 0.25) +
  geom_line(size = 1, color = "black") +
  geom_hline(yintercept = 0, linetype = "dashed", color = "gray30") +
  scale_fill_manual(name = "CI Type", breaks = c("Bootstrap", "SE"),
                    values = c("blue", "orange")) +
  xlab("Elevation (SD)") +
  ylab("log-RSS vs Mean Elevation") +
  theme_bw()
```

#### Interpreting Movement Parameters

Previously, we viewed tentative estimates of parameters describing the distribution of step-lengths and turn-angles. When estimating these parameters, however, we did not account or adjust for the influence of habitat selection. Including movement characteristics (`sl_`, `log_sl_`, and `cos_ta_`) in our iSSF allows us to update our estimates of these parameters, which describe selection-free movement distributions. Mathematical formulas required to perform these updates are given in Supplementary Appendix C. Here, we will see how to update the distributions using the functions from `amt`.

##### Update Step-Length Distribution

In our model, we included parameters for the step length (`sl_`) and its natural logarithm (`log_sl_`). We included these parameters to update the scale and shape parameters of our tentative gamma distribution, respectively. Different terms will need to be included in the iSSF, depending on the assumed step-length distribution (for details, see Supplementary Appendix C).

For our basic model – where the movement parameters are *not* interacting with any covariates – `amt` has a simple function to update the step-length distribution.

```
# Update the distribution
updated_sl <- update_sl_distr(m1)

# Check the class
class(updated_sl)
```

```
## [1] "gamma_distr" "sl_distr"    "amt_distr"   "list"
```

```
# Print
print(updated_sl)
```

```
## $name
## [1] "gamma"
## 
## $params
## $params$shape
## [1] 1.44
## 
## $params$scale
## [1] 37
## 
## 
## attr(,"class")
## [1] "gamma_distr" "sl_distr"    "amt_distr"   "list"
```

We can see that the updated distribution object has classes that are designed for working with step-length/turn-angle distributions in `amt`. In the most general sense, it is a `list`, so you can access its named components either with `$name` or `[["name"]]`, as you would with any other list. But it is also of class `amt_distr`, a general class for working with distributions in `amt`; class `sl_distr`, a more specific class for working with step-length distributions; and class `gamma_distr`, an even more specific class for working with gamma distributions. We can see that the object has a name (`"gamma"`) and the appropriate parameters for the gamma distribution – shape (`$params$shape`) and scale (`$params$scale`).

We can compare the tentative and updated distributions to see how accounting for habitat-selection changes our estimates of the selection-free step-length distribution.

```
# data.frame for plotting
plot_sl <- data.frame(x = rep(NA, 100))

# x-axis is sequence of possible step lengths
plot_sl$x <- seq(from = 0, to = 400, length.out = 100)

# y-axis is the probability density under the given gamma distribution
# For the tentative distribution
plot_sl$tentative <- dgamma(
  x = plot_sl$x, 
  shape = m1$sl_$params$shape,
  scale = m1$sl_$params$scale)

# For the updated distribution
plot_sl$updated <- dgamma(
  x = plot_sl$x,
  shape = updated_sl$params$shape,
  scale = updated_sl$params$scale)

# Pivot from wide data to long data
plot_sl <- plot_sl %>% 
  pivot_longer(cols = -x)

# Plot
ggplot(plot_sl, aes(x = x, y = value, color = factor(name))) +
  geom_line(size = 1) +
  xlab("Step Length (m)") +
  ylab("Probability Density") +
  scale_color_manual(name = "Distribution", 
                     breaks = c("tentative", "updated"),
                     values = c("blue", "orange")) +
  theme_bw()
```

In this case, our estimates do not change very much. We might have anticipated this result because the coefficients associated with `sl_` and `log_sl_` were not statistically significant in our fitted model (`m1`).

##### Update Turn-Angle Distribution

In our model, we included cosine of the turn angle (`cos_ta_`) to update the concentration parameter of our tentative von Mises distribution. Again, see Supplementary Appendix C for details of updating turn-angle distributions. Here, we will see how to use `amt` to update our tentative parameters and retrieve the selection-free turn-angle distribution.

We can use the `update_ta_distr` function in `amt` to update our turn-angle distribution:

```
# Update the distribution
updated_ta <- update_ta_distr(m1)

# Check the class
class(updated_ta)
```

```
## [1] "vonmises_distr" "ta_distr"       "amt_distr"      "list"
```

```
# Print
print(updated_ta)
```

```
## $name
## [1] "vonmises"
## 
## $params
## $params$kappa
## [1] 0.493
## 
## $params$mu
## [1] 0
## 
## 
## attr(,"class")
## [1] "vonmises_distr" "ta_distr"       "amt_distr"      "list"
```

Again, we see that the updated distribution object has several classes associated with it; it is a `list` and also of class `amt_distr`. This time it is also of class `ta_distr`, indicating that it is a turn-angle distribution, and more specifically it is a `vonmises_distr`. We can also access the appropriate parameters for the von Mises distribution – concentration (`$params$kappa`) and mean (`$params$mu` – often fixed to 0 under the assumption that animals are no more likely to turn left than right).

We can again compare the tentative and updated distributions to see how accounting for habitat-selection changes our estimates of the selection-free turn-angle distribution.

```
# data.frame for plotting
plot_ta <- data.frame(x = rep(NA, 100))

# x-axis is sequence of possible step lengths
plot_ta$x <- seq(from = -1 * pi, to = pi, length.out = 100)

# y-axis is the probability density under the given von Mises distribution
# For the tentative distribution
plot_ta$tentative <- circular::dvonmises(
  x = plot_ta$x, 
  mu = m1$ta_$params$mu,
  kappa = m1$ta_$params$kappa)

# For the updated distribution
plot_ta$updated <- circular::dvonmises(
  x = plot_ta$x, 
  mu = updated_ta$params$mu,
  kappa = updated_ta$params$kappa)

# Pivot from wide data to long data
plot_ta <- plot_ta %>% 
  pivot_longer(cols = -x)

# Plot
ggplot(plot_ta, aes(x = x, y = value, color = factor(name))) +
  geom_line(size = 1) +
  coord_cartesian(ylim = c(0, 0.25)) +
  xlab("Relative Turn Angle (radians)") +
  ylab("Probability Density") +
  scale_x_continuous(breaks = c(-pi, -pi/2, 0, pi/2, pi),
                     labels = c(expression(-pi, -pi/2, 0, pi/2, pi))) +
  scale_color_manual(name = "Distribution", 
                     breaks = c("tentative", "updated"),
                     values = c("blue", "orange")) +
  theme_bw()
```

Again, we see that our estimates do not change very much, which is not surprising given that the coefficient associated with `cos_ta_` was not statistically significant in our fitted model (`m1`).

##### Calculating RSS for Movements

Suppose we are interested in how likely our fisher is to take a 50 m step versus a 100 m step. We can propose two hypothetical locations, \(s\_1\) and \(s\_2\), which have exactly the same habitat attributes but differ in their distance from the fisher’s starting location.

We do not want to use `log_rss()` for this purpose, because the fitted movement parameters in our iSSF represent updates to the tentative step-length distribution and not the distribution itself. Instead, we want to use the updated PDF of the step-length distribution directly. We can calculate the likelihood under the selection-free step-length distribution of taking a 50 m step and the likelihood of taking a 100 m step. Then we divide for an estimate of how much more likely a 50 m step is than a 100 m step.

```
like50 <- dgamma(50, 
                 shape = updated_sl$params$shape,
                 scale = updated_sl$params$scale)
like100 <- dgamma(100, 
                  shape = updated_sl$params$shape,
                  scale = updated_sl$params$scale)
like50/like100
```

```
## [1] 2.85
```

Thus, we conclude Lupe is nearly 3 times more likely to take the shorter (50 m versus 100 m) step.

### More Complex Models

#### Non-Linear Relationships: Adding Quadratic Terms

Suppose, now, that we hypothesize that our fisher selects for intermediate elevations and avoids locations with extremely low or high elevations. We could use splines or polynomials to capture this non-linear effect. Here, we consider adding a quadratic term for elevation in our model.

```
m2 <- ssf_dat %>% 
  fit_issf(case_ ~ popden_end + elevation_end + I(elevation_end^2) + 
             landuseC_end + sl_ + log_sl_ + cos_ta_ + 
             strata(step_id_), model = TRUE)
```

In our basic model, we could interpret the fitted \(\beta\_{elevation}\) as capturing the log relative selection strength (or log-RSS) associated with a one unit change in elevation. With the addition of the quadratic term, the log-RSS for one unit change in elevation is no longer constant.  To illustrate this point, we compare the log-RSS for a one unit change in elevation for our two models (`m1` has just a linear term for elevation; `m2` has a quadratic term for elevation), using two different reference values for `elevation`:

- \(s\_1\): `elevation_end = 3`, `popden_end = 1.5`, `landuseC_end = "wet"`
- \(s\_2\): `elevation_end = 2`, `popden_end = 1.5`, `landuseC_end = "wet"`
- \(s\_3\): `elevation_end = -1`, `popden_end = 1.5`, `landuseC_end = "wet"`
- \(s\_4\): `elevation_end = -2`, `popden_end = 1.5`, `landuseC_end = "wet"`

We can see that \(s\_1\) and \(s\_2\) are separated by 1 unit of elevation, and \(s\_3\) and \(s\_4\) are also separated by 1 unit of elevation. Let’s compare the two sets of locations.

```
# data.frame for s1
s1 <- data.frame(
  elevation_end = 3, # mean of elev, since we scaled and centered
  popden_end = 1.5,
  landuseC_end = factor("wet", levels = levels(ssf_dat$landuseC_end)),
  sl_ = 100,
  log_sl_ = log(100),
  cos_ta_ = 1)

# data.frame for s2
s2 <- data.frame(
  elevation_end = 2, # mean of elev, since we scaled and centered
  popden_end = 1.5,
  landuseC_end = factor("wet", levels = levels(ssf_dat$landuseC_end)),
  sl_ = 100,
  log_sl_ = log(100),
  cos_ta_ = 1)

# data.frame for s3
s3 <- data.frame(
  elevation_end = -1, # mean of elev, since we scaled and centered
  popden_end = 1.5,
  landuseC_end = factor("wet", levels = levels(ssf_dat$landuseC_end)),
  sl_ = 100,
  log_sl_ = log(100),
  cos_ta_ = 1)

# data.frame for s4
s4 <- data.frame(
  elevation_end = -2, # mean of elev, since we scaled and centered
  popden_end = 1.5,
  landuseC_end = factor("wet",  levels = levels(ssf_dat$landuseC_end)),
  sl_ = 100,
  log_sl_ = log(100),
  cos_ta_ = 1)

# Calculate log-RSS under m1
lr_m1_s1s2 <- log_rss(m1, s1, s2)
lr_m1_s3s4 <- log_rss(m1, s3, s4)

# Calculate log-RSS under m2
lr_m2_s1s2 <- log_rss(m2, s1, s2)
lr_m2_s3s4 <- log_rss(m2, s3, s4)

# Compare
data.frame(
  "model" = c("linear", "quadratic"),
  "logRSS_s1_s2" = round(c(lr_m1_s1s2$df$log_rss, lr_m2_s1s2$df$log_rss), 3),
  "logRSS_s3_s4" = round(c(lr_m1_s3s4$df$log_rss, lr_m2_s3s4$df$log_rss), 3))
```

```
##       model logRSS_s1_s2 logRSS_s3_s4
## 1    linear        0.170        0.170
## 2 quadratic        0.047        0.269
```

Notice that the reference value for `elevation_end` (`x1`) does not impact the log-RSS associated with a 1 unit change when the model is linear. If the model includes the quadratic term, however, then the reference `elevation` value is important.

If we plot a range of log-RSS predictions, we can see the slope slowly leveling off as the value of `elevation` increases.

```
# Make a new data.frame for s1
s1 <- data.frame(
  elevation_end = seq(from = -2, to = 2, length.out = 100),
  popden_end = 1.5,
  landuseC_end = factor("wet", levels = levels(ssf_dat$landuseC_end)),
  sl_ = 100,
  log_sl_ = log(100),
  cos_ta_ = 1)

# data.frame for s2
s2 <- data.frame(
  elevation_end = 0, # mean of elev, since we scaled and centered
  popden_end = 1.5,
  landuseC_end = factor("wet", levels = levels(ssf_dat$landuseC_end)),
  sl_ = 100,
  log_sl_ = log(100),
  cos_ta_ = 1)

# Calculate log-RSS
lr_m2 <- log_rss(m2, s1, s2)

# Plot using ggplot2
ggplot(lr_m2$df, aes(x = elevation_end_x1, y = log_rss)) +
  geom_line(size = 1) +
  geom_hline(yintercept = 0, linetype = "dashed", color = "gray30") +
  xlab("Elevation (SD)") +
  ylab("log-RSS vs Mean Elevation") +
  theme_bw()
```

Our hypothesis that Lupe selects intermediate elevations doesn’t seem to have support here (at least over the range of elevations within Lupe’s MCP), and the non-linear effect that is estimated is rather muted. Nevertheless, quadratic terms are often useful for predictors that may represent *conditions* (see main text; Matthiopoulos et al., 2015).

#### Movement and Habitat Interactions

In all of our previous models, we have assumed that a single step-length distribution and a single turn-angle distribution was sufficient for modeling selection-free movements. However, considering interactions between movement properties can provide valuable insight into an animal’s motivations for moving (e.g., Dickie, McNay, Sutherland, Cody, & Avgar, 2020). In this section, we will demonstrate how to fit and interpret models that include interactions between movement characteristics and categorical or continuous explanatory variables capturing environmental characteristics.

##### Interactions Between Movement Characteristics and Categorical Variables

Suppose we hypothesize that our fisher moves differently depending on the land-use class that it is in at the start of the movement step. We can fit a model where all of the movement characteristics (`sl_`, `log_sl_`, an d`cos_ta_`) interact with `landuseC` as follows:

```
m3 <- ssf_dat %>% 
  fit_issf(case_ ~ popden_end + elevation_end +  
             landuseC_end + 
             sl_ + log_sl_ + cos_ta_ + 
             landuseC_start:(sl_ + log_sl_ + cos_ta_) +
             strata(step_id_), model = TRUE)
```

Under this model, Lupe’s selection-free movement kernel, \(\phi(s,s';\gamma(\Delta t,t,X(s,t)))\), depends on the value of `landuseC` at the start of the movement step and Lupe’s step-selection function, \(w(X(s,t+\Delta t);\beta(\Delta t))\), depends on the value of`landuseC` at the end of the movement step.

Let’s look at how the step-length distributions vary depending on the land-use class. Earlier, we saw simple `amt` functions to update step-length and turn-angle distributions from a fitted model. However, these easy-to-use functions cannot accommodate more complex models that include interactions between habitat covariates and movement characteristics. Instead, we will have to use a separate set of functions that require manual specification of the appropriate \(\beta\)’s (see `?update_distr_man`).

Before we demonstrate these functions, let’s take a look at the parameters in the fitted model. For updating the step-length distribution, pay attention to those parameters that interact with `sl_` or `log_sl_`.

```
summary(m3)
```

```
## Call:
## coxph(formula = Surv(rep(1, 16175L), case_) ~ popden_end + elevation_end + 
##     landuseC_end + sl_ + log_sl_ + cos_ta_ + landuseC_start:(sl_ + 
##     log_sl_ + cos_ta_) + strata(step_id_), data = data, model = TRUE, 
##     method = "exact")
## 
##   n= 16175, number of events= 1155 
## 
##                                  coef exp(coef)  se(coef)     z Pr(>|z|)   
## popden_end                  -0.107154  0.898387  0.106971 -1.00   0.3165   
## elevation_end                0.173319  1.189245  0.060050  2.89   0.0039 **
## landuseC_endgrass           -0.851573  0.426743  0.519500 -1.64   0.1012   
## landuseC_endwet             -0.242487  0.784674  0.420745 -0.58   0.5644   
## sl_                          0.000659  1.000660  0.001337  0.49   0.6220   
## log_sl_                      0.034202  1.034794  0.059130  0.58   0.5630   
## cos_ta_                      0.020188  1.020393  0.046737  0.43   0.6658   
## sl_:landuseC_startgrass     -0.047730  0.953391  0.045496 -1.05   0.2941   
## sl_:landuseC_startwet        0.002498  1.002501  0.007282  0.34   0.7316   
## log_sl_:landuseC_startgrass  3.935494 51.187450  2.998270  1.31   0.1893   
## log_sl_:landuseC_startwet   -0.346018  0.707500  0.271696 -1.27   0.2028   
## cos_ta_:landuseC_startgrass  0.462415  1.587904  0.758162  0.61   0.5419   
## cos_ta_:landuseC_startwet   -0.295292  0.744314  0.213181 -1.39   0.1660   
## ---
## Signif. codes:  0 '***' 0.001 '**' 0.01 '*' 0.05 '.' 0.1 ' ' 1
## 
##                             exp(coef) exp(-coef) lower .95 upper .95
## popden_end                      0.898     1.1131     0.728      1.11
## elevation_end                   1.189     0.8409     1.057      1.34
## landuseC_endgrass               0.427     2.3433     0.154      1.18
## landuseC_endwet                 0.785     1.2744     0.344      1.79
## sl_                             1.001     0.9993     0.998      1.00
## log_sl_                         1.035     0.9664     0.922      1.16
## cos_ta_                         1.020     0.9800     0.931      1.12
## sl_:landuseC_startgrass         0.953     1.0489     0.872      1.04
## sl_:landuseC_startwet           1.003     0.9975     0.988      1.02
## log_sl_:landuseC_startgrass    51.187     0.0195     0.144  18251.37
## log_sl_:landuseC_startwet       0.708     1.4134     0.415      1.21
## cos_ta_:landuseC_startgrass     1.588     0.6298     0.359      7.02
## cos_ta_:landuseC_startwet       0.744     1.3435     0.490      1.13
## 
## Concordance= 0.526  (se = 0.011 )
## Likelihood ratio test= 29.2  on 13 df,   p=0.006
## Wald test            = 23.1  on 13 df,   p=0.04
## Score (logrank) test = 26.6  on 13 df,   p=0.01
```

Recall that one of the `landuseC` categories is treated as a reference category (in this case, `"forest"`). We update the parameters of the step-length distribution for the reference category with the main effect parameters (i.e., `sl_` and `log_sl_`). For the other categories, we need to add the appropriate interaction terms to these parameters.

```
# Forest step-length distribution
forest_sl <- update_gamma(
  dist = m3$sl_,
  beta_sl = m3$model$coefficients["sl_"],
  beta_log_sl = m3$model$coefficients["log_sl_"])

# Grass step-length distribution
grass_sl <- update_gamma(
  dist = m3$sl_,
  beta_sl = m3$model$coefficients["sl_"] +
    m3$model$coefficients["sl_:landuseC_startgrass"],
  beta_log_sl = m3$model$coefficients["log_sl_"] +
    m3$model$coefficients["log_sl_:landuseC_startgrass"])

# Wet step-length distribution
wet_sl <- update_gamma(
  dist = m3$sl_,
  beta_sl = m3$model$coefficients["sl_"] +
    m3$model$coefficients["sl_:landuseC_startwet"],
  beta_log_sl = m3$model$coefficients["log_sl_"] + 
    m3$model$coefficients["log_sl_:landuseC_startwet"])
```

We can follow a similar process with the turn-angle distribution.

```
# Forest turn-angle distribution
forest_ta <- update_vonmises(
  dist = m3$ta_, beta_cos_ta = m3$model$coefficients["cos_ta_"])

# Grass turn-angle distribution
grass_ta <- update_vonmises(
  dist = m3$ta_, 
  beta_cos_ta = m3$model$coefficients["cos_ta_"] +
    m3$model$coefficients["cos_ta_:landuseC_startgrass"])

# Wet turn-angle distribution
wet_ta <- update_vonmises(
  dist = m3$ta_,
  beta_cos_ta = m3$model$coefficients["cos_ta_"] +
    m3$model$coefficients["cos_ta_:landuseC_startwet"])
```

Now, we can plot the original and updated distributions for each `landuseC`.

```
# data.frame for plotting
plot_sl <- data.frame(x = rep(NA, 100))

# x-axis is sequence of possible step lengths
plot_sl$x <- seq(from = 0, to = 400, length.out = 100)

# y-axis is the probability density under the given gamma distribution
# Forest
plot_sl$forest <- dgamma(x = plot_sl$x, 
                         shape = forest_sl$params$shape,
                         scale = forest_sl$params$scale)
# Grass
plot_sl$grass <- dgamma(x = plot_sl$x, 
                        shape = grass_sl$params$shape,
                        scale = grass_sl$params$scale)
# Wet
plot_sl$wet <- dgamma(x = plot_sl$x, 
                      shape = wet_sl$params$shape,
                      scale = wet_sl$params$scale)

# Pivot from wide to long data
plot_sl <- plot_sl %>% 
  pivot_longer(cols = -x)

# Plot
p1<-ggplot(plot_sl, aes(x = x, y = value, color = factor(name))) +
  geom_line(size = 1) +
  scale_color_manual(name = "Land-use",
                       breaks = c("forest", "grass", "wet"),
                       values = c("forestgreen", "wheat", "blue")) +
  xlab("Step Length (m)") +
  ylab("Probability Density") +
  theme_bw()
 

#data.frame for plotting
plot_ta <- data.frame(x = rep(NA, 100))

# x-axis is sequence of possible step lengths
plot_ta$x <- seq(from = -pi, to = pi, length.out = 100)

# y-axis is the probability density under the given gamma distribution
# Forest
plot_ta$forest <- circular::dvonmises(x = plot_ta$x, 
                      kappa = forest_ta$params$kappa,
                      mu = 0)
# Grass
plot_ta$grass <- circular::dvonmises(x = plot_ta$x, 
                      kappa = grass_ta$params$kappa,
                      mu = 0)
# Wet
plot_ta$wet <- circular::dvonmises(x = plot_ta$x, 
                      kappa = wet_ta$params$kappa,
                      mu = 0)

# Pivot from wide to long data
plot_ta <- plot_ta %>% 
  pivot_longer(cols = -x)

# Plot
p2 <- ggplot(plot_ta, aes(x = x, y = value, color = factor(name))) +
  geom_line(size = 1) +
  scale_color_manual(name = "Land-use",
                     breaks = c("forest", "grass", "wet"),
                     values = c("forestgreen", "wheat", "blue")) +
  scale_x_continuous(breaks = c(-pi, -pi/2, 0, pi/2, pi),
                     labels = c(expression(-pi, -pi/2, 0, pi/2, pi))) +
  xlab("Turn Angle (radians)") +
  ylab("Probability Density") +
  theme_bw()

combined <- p1 + p2 & theme(legend.position = "bottom")
combined + plot_layout(guides = "collect")
```

We see that Lupe tends to take larger, more directed steps when in `grass` and slower and more tortuous steps in `wet` habitat. One possibility is that Lupe tends to travel through `grass` and forage in `forest` and `wet` habitats.

##### Continuous Movement Predictors

We can follow a similar process for continuous movement predictors. Suppose we want to know if Lupe moves differently depending on the elevation she finds herself in. We can fit the following model:

```
m4 <- ssf_dat %>% 
  fit_issf(case_ ~ popden_end + landuseC_end +   
             elevation_end + 
             sl_ + log_sl_ + cos_ta_ + 
             elevation_start:(sl_ + log_sl_ + cos_ta_) + 
             strata(step_id_), model = TRUE)
```

Again, let’s inspect the fitted model:

```
summary(m4)
```

```
## Call:
## coxph(formula = Surv(rep(1, 16175L), case_) ~ popden_end + landuseC_end + 
##     elevation_end + sl_ + log_sl_ + cos_ta_ + elevation_start:(sl_ + 
##     log_sl_ + cos_ta_) + strata(step_id_), data = data, model = TRUE, 
##     method = "exact")
## 
##   n= 16175, number of events= 1155 
## 
##                              coef exp(coef)  se(coef)     z Pr(>|z|)  
## popden_end              -0.109319  0.896444  0.107207 -1.02    0.308  
## landuseC_endgrass       -1.199118  0.301460  0.511767 -2.34    0.019 *
## landuseC_endwet         -0.015773  0.984350  0.369561 -0.04    0.966  
## elevation_end            0.157844  1.170984  0.062294  2.53    0.011 *
## sl_                      0.000765  1.000765  0.001309  0.58    0.559  
## log_sl_                  0.025121  1.025439  0.057641  0.44    0.663  
## cos_ta_                  0.012051  1.012124  0.045574  0.26    0.791  
## sl_:elevation_start      0.000828  1.000828  0.001332  0.62    0.534  
## log_sl_:elevation_start -0.066514  0.935650  0.058371 -1.14    0.254  
## cos_ta_:elevation_start -0.097660  0.906958  0.045985 -2.12    0.034 *
## ---
## Signif. codes:  0 '***' 0.001 '**' 0.01 '*' 0.05 '.' 0.1 ' ' 1
## 
##                         exp(coef) exp(-coef) lower .95 upper .95
## popden_end                  0.896      1.116     0.727     1.106
## landuseC_endgrass           0.301      3.317     0.111     0.822
## landuseC_endwet             0.984      1.016     0.477     2.031
## elevation_end               1.171      0.854     1.036     1.323
## sl_                         1.001      0.999     0.998     1.003
## log_sl_                     1.025      0.975     0.916     1.148
## cos_ta_                     1.012      0.988     0.926     1.107
## sl_:elevation_start         1.001      0.999     0.998     1.003
## log_sl_:elevation_start     0.936      1.069     0.835     1.049
## cos_ta_:elevation_start     0.907      1.103     0.829     0.992
## 
## Concordance= 0.537  (se = 0.011 )
## Likelihood ratio test= 24.6  on 10 df,   p=0.006
## Wald test            = 23.8  on 10 df,   p=0.008
## Score (logrank) test = 24.1  on 10 df,   p=0.007
```

We can see that the interactions between the step-length parameters and `elevation` are not detectable, so we shouldn’t expect much of an effect of `elevation` on the step-length distribution. However, the relationship with the turn-angle parameter (`cos_ta_`) is stronger, so we might see an effect there. Let’s update our step-length and turn-angle distributions and see how they look.

Because we have included interactions between movement characteristics and a continuous covariate, `elevation`, our model allows for an infinite number of selection-free movement kernels – i.e., Lupe’s selection-free movement kernel will depend on the specific value of `elevation` associated with her current location. For example, the scale parameter of the gamma distribution, \(\beta\_{l}\), is a function of elevation:

\[\beta\_l = \beta\_{sl\\_} + \beta\_{elevation:sl\\_} \times elevation\]

To help visualize the effect of `elevation` on movement, it can be helpful to plot selection-free movement kernels for a range of `elevation` values. Below, we pick three values of `elevation` (-2, 0, 2) which we label (“low”, “medium”, and “high”), respectively. Remember that we scaled and centered our covariates so these values correspond to the mean and \(\pm 2\) standard deviations away from the mean. We’ll create an updated gamma step-length distribution for each of these three values of `elevation`:

```
# Low elevation step-length distribution
low_sl <- update_gamma(
  dist = m4$sl_,
  beta_sl = m4$model$coefficients["sl_"] +
    -2 * m4$model$coefficients["sl_:elevation_start"],
  beta_log_sl = m4$model$coefficients["log_sl_"] +
    -2 * m4$model$coefficients["log_sl_:elevation_start"])

# Medium elevation step-length distribution
med_sl <- update_gamma(
  dist = m4$sl_,
  beta_sl = m4$model$coefficients["sl_"] +
    0 * m4$model$coefficients["sl_:elevation_start"],
  beta_log_sl = m4$model$coefficients["log_sl_"] +
    0 * m4$model$coefficients["log_sl_:elevation_start"])

# Wet step-length distribution
hi_sl <- update_gamma(
  dist = m4$sl_,
  beta_sl = m4$model$coefficients["sl_"] +
    2 * m4$model$coefficients["sl_:elevation_start"],
  beta_log_sl = m4$model$coefficients["log_sl_"] +
    2 * m4$model$coefficients["log_sl_:elevation_start"])
```

Similarly, we calculate updated turn-angle distributions for each of these 3 values of `elevation`.

```
# low turn-angle distribution
low_ta <- update_vonmises(
  dist = m4$ta_,
  beta_cos_ta = m4$model$coefficients["cos_ta_"] +
    -2 * m4$model$coefficients["cos_ta_:elevation_start"])

# med turn-angle distribution
med_ta <- update_vonmises(
  dist = m4$ta_,
  beta_cos_ta = m4$model$coefficients["cos_ta_"] +
    0 * m4$model$coefficients["cos_ta_:elevation_start"])

# hi turn-angle distribution
hi_ta <- update_vonmises(
  dist = m4$ta_,
  beta_cos_ta = m4$model$coefficients["cos_ta_"] +
    2 * m4$model$coefficients["cos_ta_:elevation_start"])
```

Now, let’s plot the results:

```
#data.frame for plotting
plot_sl <- data.frame(x = rep(NA, 100))

# x-axis is sequence of possible step lengths
plot_sl$x <- seq(from = 0, to = 400, length.out = 100)

# y-axis is the probability density under the given gamma distribution
# Forest
plot_sl$low <- dgamma(x = plot_sl$x, 
                      shape = low_sl$params$shape,
                      scale = low_sl$params$scale)
# Grass
plot_sl$medium <- dgamma(x = plot_sl$x, 
                         shape = med_sl$params$shape,
                         scale = med_sl$params$scale)
# Wet
plot_sl$high <- dgamma(x = plot_sl$x, 
                       shape = hi_sl$params$shape,
                       scale = hi_sl$params$scale)

# Pivot from wide to long data
plot_sl <- plot_sl %>% 
  pivot_longer(cols = -x)

# Plot
p1 <- ggplot(plot_sl, aes(x = x, y = value, color = factor(name))) +
  geom_line(size = 1) +
  scale_color_manual(name = "Elevation",
                     breaks = c("low", "medium", "high"),
                     values = c("navyblue", "gray50", "firebrick")) +
  xlab("Step Length (m)") +
  ylab("Probability Density") +
  theme_bw() 

#data.frame for plotting
plot_ta <- data.frame(x = rep(NA, 100))

# x-axis is sequence of possible step lengths
plot_ta$x <- seq(from = -pi, to = pi, length.out = 100)

# y-axis is the probability density under the given gamma distribution
# low
plot_ta$low <- circular::dvonmises(x = plot_ta$x, 
                                   kappa = low_ta$params$kappa,
                                   mu = 0)
# med
plot_ta$medium <- circular::dvonmises(x = plot_ta$x, 
                      kappa = med_ta$params$kappa,
                      mu = 0)
# hi
plot_ta$high <- circular::dvonmises(x = plot_ta$x, 
                      kappa = hi_ta$params$kappa,
                      mu = 0)

# Pivot from wide to long data
plot_ta <- plot_ta %>% 
  pivot_longer(cols = -x)

# Plot
p2 <- ggplot(plot_ta, aes(x = x, y = value, color = factor(name))) +
  geom_line(size = 1) +
  scale_color_manual(name = "Elevation",
                     breaks = c("low", "medium", "high"),
                     values = c("navyblue", "gray50", "firebrick")) +
  scale_x_continuous(breaks = c(-pi, -pi/2, 0, pi/2, pi),
                     labels = c(expression(-pi, -pi/2, 0, pi/2, pi))) +
  xlab("Turn Angle (radians)") +
  ylab("Probability Density") +
  theme_bw()
combined <- p1 + p2 & theme(legend.position = "bottom")
combined + plot_layout(guides = "collect")
```

The step-length distributions are very similar, which we expected based on the non-significant coefficients associated with the interactions between `elevation` and (`sl_` and `log_sl_`) in our fitted model. We see bigger differences in the turn-angle distributions, as we expected based on the statistical significance of the `elevation` and `cos_ta_` interactions. Our fisher has more directed movements at low elevation and less directed movements at high elevation.

### Conclusion

We have demonstrated how we can fit a rich set of models that capture the effects of environmental predictors on animal movement and habitat selection using the `amt` package in R.

### Session

```
sessioninfo::session_info()
```

```
## - Session info ---------------------------------------------------------------
##  setting  value                       
##  version  R version 4.0.2 (2020-06-22)
##  os       Windows 10 x64              
##  system   x86_64, mingw32             
##  ui       RTerm                       
##  language (EN)                        
##  collate  English_United States.1252  
##  ctype    English_United States.1252  
##  tz       America/Chicago             
##  date     2021-01-22                  
## 
## - Packages -------------------------------------------------------------------
##  package      * version  date       lib source                       
##  amt          * 0.1.3    2020-11-04 [1] Github (jmsigner/amt@ee58ca1)
##  assertthat     0.2.1    2019-03-21 [1] CRAN (R 4.0.2)               
##  backports      1.1.7    2020-05-13 [1] CRAN (R 4.0.0)               
##  bibtex         0.4.2.2  2020-01-02 [1] CRAN (R 4.0.2)               
##  blob           1.2.1    2020-01-20 [1] CRAN (R 4.0.2)               
##  boot           1.3-25   2020-04-26 [2] CRAN (R 4.0.2)               
##  broom        * 0.7.0    2020-07-09 [1] CRAN (R 4.0.2)               
##  cellranger     1.1.0    2016-07-27 [1] CRAN (R 4.0.2)               
##  checkmate      2.0.0    2020-02-06 [1] CRAN (R 4.0.2)               
##  circular       0.4-93   2017-06-29 [1] CRAN (R 4.0.2)               
##  cli            2.0.2    2020-02-28 [1] CRAN (R 4.0.2)               
##  codetools      0.2-16   2018-12-24 [2] CRAN (R 4.0.2)               
##  colorspace     1.4-1    2019-03-18 [1] CRAN (R 4.0.2)               
##  crayon         1.3.4    2017-09-16 [1] CRAN (R 4.0.2)               
##  DBI            1.1.0    2019-12-15 [1] CRAN (R 4.0.2)               
##  dbplyr         1.4.4    2020-05-27 [1] CRAN (R 4.0.2)               
##  digest         0.6.25   2020-02-23 [1] CRAN (R 4.0.2)               
##  dplyr        * 1.0.1    2020-07-31 [1] CRAN (R 4.0.2)               
##  ellipsis       0.3.1    2020-05-15 [1] CRAN (R 4.0.2)               
##  evaluate       0.14     2019-05-28 [1] CRAN (R 4.0.2)               
##  fansi          0.4.1    2020-01-08 [1] CRAN (R 4.0.2)               
##  farver         2.0.3    2020-01-16 [1] CRAN (R 4.0.2)               
##  fitdistrplus   1.1-1    2020-05-19 [1] CRAN (R 4.0.2)               
##  forcats      * 0.5.0    2020-03-01 [1] CRAN (R 4.0.2)               
##  fs             1.5.0    2020-07-31 [1] CRAN (R 4.0.2)               
##  gbRd           0.4-11   2012-10-01 [1] CRAN (R 4.0.0)               
##  generics       0.0.2    2018-11-29 [1] CRAN (R 4.0.2)               
##  ggplot2      * 3.3.2    2020-06-19 [1] CRAN (R 4.0.2)               
##  glue           1.4.1    2020-05-13 [1] CRAN (R 4.0.2)               
##  gtable         0.3.0    2019-03-25 [1] CRAN (R 4.0.2)               
##  haven          2.3.1    2020-06-01 [1] CRAN (R 4.0.2)               
##  hms            0.5.3    2020-01-08 [1] CRAN (R 4.0.2)               
##  htmltools      0.5.0    2020-06-16 [1] CRAN (R 4.0.2)               
##  httr           1.4.2    2020-07-20 [1] CRAN (R 4.0.2)               
##  jsonlite       1.7.0    2020-06-25 [1] CRAN (R 4.0.2)               
##  knitr        * 1.30     2020-09-22 [1] CRAN (R 4.0.3)               
##  labeling       0.3      2014-08-23 [1] CRAN (R 4.0.0)               
##  lattice        0.20-41  2020-04-02 [2] CRAN (R 4.0.2)               
##  lifecycle      0.2.0    2020-03-06 [1] CRAN (R 4.0.2)               
##  lubridate      1.7.9    2020-06-08 [1] CRAN (R 4.0.2)               
##  magrittr       1.5      2014-11-22 [1] CRAN (R 4.0.2)               
##  MASS           7.3-51.6 2020-04-26 [2] CRAN (R 4.0.2)               
##  Matrix         1.2-18   2019-11-27 [1] CRAN (R 4.0.2)               
##  modelr         0.1.8    2020-05-19 [1] CRAN (R 4.0.2)               
##  munsell        0.5.0    2018-06-12 [1] CRAN (R 4.0.2)               
##  mvtnorm        1.1-1    2020-06-09 [1] CRAN (R 4.0.0)               
##  patchwork    * 1.0.1    2020-06-22 [1] CRAN (R 4.0.2)               
##  pillar         1.4.6    2020-07-10 [1] CRAN (R 4.0.2)               
##  pkgconfig      2.0.3    2019-09-22 [1] CRAN (R 4.0.2)               
##  purrr        * 0.3.4    2020-04-17 [1] CRAN (R 4.0.2)               
##  R6             2.4.1    2019-11-12 [1] CRAN (R 4.0.2)               
##  raster         3.3-13   2020-07-17 [1] CRAN (R 4.0.2)               
##  Rcpp           1.0.5    2020-07-06 [1] CRAN (R 4.0.2)               
##  Rdpack         1.0.0    2020-07-01 [1] CRAN (R 4.0.2)               
##  readr        * 1.3.1    2018-12-21 [1] CRAN (R 4.0.2)               
##  readxl         1.3.1    2019-03-13 [1] CRAN (R 4.0.2)               
##  reprex         0.3.0    2019-05-16 [1] CRAN (R 4.0.2)               
##  rlang          0.4.7    2020-07-09 [1] CRAN (R 4.0.2)               
##  rmarkdown      2.3      2020-06-18 [1] CRAN (R 4.0.2)               
##  rstudioapi     0.11     2020-02-07 [1] CRAN (R 4.0.2)               
##  rvest          0.3.6    2020-07-25 [1] CRAN (R 4.0.2)               
##  scales         1.1.1    2020-05-11 [1] CRAN (R 4.0.2)               
##  sessioninfo    1.1.1    2018-11-05 [1] CRAN (R 4.0.2)               
##  sp             1.4-2    2020-05-20 [1] CRAN (R 4.0.2)               
##  stringi        1.4.6    2020-02-17 [1] CRAN (R 4.0.0)               
##  stringr      * 1.4.0    2019-02-10 [1] CRAN (R 4.0.2)               
##  survival       3.1-12   2020-04-10 [2] CRAN (R 4.0.2)               
##  tibble       * 3.0.3    2020-07-10 [1] CRAN (R 4.0.2)               
##  tidyr        * 1.1.1    2020-07-31 [1] CRAN (R 4.0.2)               
##  tidyselect     1.1.0    2020-05-11 [1] CRAN (R 4.0.2)               
##  tidyverse    * 1.3.0    2019-11-21 [1] CRAN (R 4.0.2)               
##  utf8           1.1.4    2018-05-24 [1] CRAN (R 4.0.2)               
##  vctrs          0.3.2    2020-07-15 [1] CRAN (R 4.0.2)               
##  withr          2.2.0    2020-04-20 [1] CRAN (R 4.0.2)               
##  xfun           0.16     2020-07-24 [1] CRAN (R 4.0.2)               
##  xml2           1.3.2    2020-04-23 [1] CRAN (R 4.0.2)               
##  yaml           2.2.1    2020-02-01 [1] CRAN (R 4.0.2)               
## 
## [1] C:/Users/jfieberg/Documents/R/win-library/4.0
## [2] C:/Program Files/R/R-4.0.2/library
```
