## Supplementary Appendix C for "A ‘How-to’ Guide for Interpreting Parameters in Habitat-Selection Analyses": AppC_iSSA_movement.html

The movement parameters that may be included in an iSSF to update the selection-free step length and turning angle distributions depend on the distribution that the user chooses to model these movement properties.

Once the user decides on the appropriate distribution, they must choose tentative parameters for that distribution. Often these are chosen by quickly fitting the parameters of the distribution to the observed step lengths/turning angles in the observed data (see Appendix B for examples).

After fitting the iSSF, the parameters of the tentative distribution are updated with the fitted values (the \(\beta\)s) from the model, resulting in the new parameters for the selection-free distribution.

### Step Length Distributions

#### Exponential

If available step lengths were sampled from an exponential distribution with tentative rate parameter \(\lambda\_0\), and the step length (\(l\)) was included as a covariate in the analysis, with resulting coefficient estimate \(\beta\_l\), the adjusted (selection-free) step length Exponential rate parameter (\(\hat{\lambda}\)) is given by:

\[\hat{\lambda} = \lambda\_0 - \beta\_l\]

#### Half-normal

If available step lengths were sampled from a half-normal distribution with scale parameter (standard deviation) \(\sigma\_0\), and the squared step length (\(l^2\)) was included as a covariate in the analysis, with resulting coefficient estimate \(\beta\_{l^2}\), the adjusted (selection-free) step length half-normal scale parameter (\(\hat{\sigma}\)) is given by:

\[\hat{\sigma} = \frac{\sigma\_0}{\sqrt{1 - 2 \sigma\_0^2 \beta\_{l^2}}}\]

#### Log-normal

If available step lengths were sampled from a log-normal distribution with tentative mean \(\mu\_0\) and standard deviation \(\sigma\_0\), and the log-transformed step length (\(ln[l]\)) and its square (\(ln[l]^2\)) were included as covariates in the analysis, with resulting coefficient estimates \(\beta\_{ln[l]}\) and \(\beta\_{ln[l]^2}\) (respectively), the adjusted (selection-free) step length log-normal mean (\(\hat{\mu}\)) and standard-deviation (\(\hat{\sigma}\)) parameters are given by:

\[
\left\{
\begin{array}{c}
\hat{\mu} = \frac{\mu\_0 - \sigma\_0\beta\_{ln[l]}}{1 - 2 \sigma\_0^2 \beta\_{ln[l]^2}} \\[2ex]
\hat{\sigma} = \frac{\sigma\_0}{\sqrt{1 - 2 \sigma\_0^2 \beta\_{ln[l]^2}}}
\end{array}
\right.
\]

#### Gamma

If available step lengths were sampled from a gamma distribution with tentative shape \(k\_0\) and scale \(q\_0\), and the step length (\(l\)) and its log-transform (\(ln[l]\)) were included as covariates in the analysis, with resulting coefficient estimates \(\beta\_l\) and \(\beta\_{ln[l]}\) (respectively), the adjusted (selection-free) step length Gamma shape (\(\hat{k}\)) and scale (\(\hat{q}\)) parameters are given by:

\[
\left\{
\begin{array}{c}
\hat{k} = k\_0 + \beta\_{ln[l]} \\[2ex]
\hat{q} = \frac{1}{\left(\frac{1}{q\_0} - \beta\_l \right)}
\end{array}
\right.
\]

### Turning Angle Distributions

#### Von Mises

If available turning angles were sampled from a von Mises distribution with tentative concentration parameter \(\nu\_0\), and the cosine of the turn angle (\(cos[\theta]\)) was included as a covariate in the analysis, with resulting coefficient estimate \(\beta\_{cos[θ]}\), the adjusted (selection-free) von Mises concentration parameter (\(\hat{\nu}\)) is given by:

\[\hat{\nu} = \nu\_0 + \beta\_{cos[\theta]}\]

#### Uniform

Similarly, if available turning angles were sampled from a Uniform distribution, that is equivalent to sampling from a von Mises distribution with tentative concentration parameter \(\nu\_0 = 0\). Thus, if the cosine of the turn angle (\(cos[\theta]\)) was included as a covariate in the analysis, with resulting coefficient estimate \(\beta\_{cos[θ]}\), the adjusted (selection-free) von Mises concentration parameter (\(\hat{\nu}\)) is given by:

\[\hat{\nu} = \nu\_0 + \beta\_{cos[\theta]} = 0 + \beta\_{cos[\theta]}\]

\[ = \beta\_{cos[\theta]}\]
